# Autonomous AI agents discover reverse transcriptases with tandem repeat arrays

**DOI:** 10.64898/2026.10.05.755809

**Authors:** Peter H. Yoon, Januka S. Athukoralage, Emmanuel Ameisen, Eric Kauderer-Abrams, Nicholas T. Perry, Matthew G. Durrant

## Abstract

Metagenomic databases hold vast quantities of unexplored genetic material that can seed tools for biotechnology and medicine, but discovery is limited by our ability to examine sequence data at scale. Here, we show that large language models (LLMs) in an autonomous harness can alleviate this bottleneck and surface novel molecular systems. We deployed Claude Code instances to survey reverse transcriptase (RT) loci across 1.9 billion protein clusters. Among the top candidates were array-associated RTs (ART), a novel family of jumbo-phage RTs coupled with an array of ∼200-nucleotide units and a dedicated partner gene. Session transcripts show that Claude identified the ART family by directly examining its DNA sequences and recognizing the atypical repeats, a behavior attributable to specific Mythos 5 internal signals that respond to repeated DNA. ART arrays are highly expressed and appear as discrete units during *Staphylococcus* phage infection, suggesting an RT system directed by a repertoire of distinct RNAs. These findings indicate that LLMs can autonomously detect anomalies and drive analyses to initiate biological discoveries.

## Introduction

Sequence databases have become the largest and fastest-growing record of biological diversity (1–4). Mining them has yielded industrial enzymes, new antibiotics, and genome-engineering tools (5–7). Many of these discoveries began with a researcher inspecting sequences and recognizing atypical features (8–10). Driven by metagenomic sequencing, these databases now double every four years and hold billions of protein sequences, increasing not only the diversity available for discovery but also the volume of data that must be examined (2, 3, 11, 12).

Systematic study of databases of this size is possible only through computational pipelines, which retrieve homologs of a known system’s components and sort the hits into classes by a set of predefined features (13). Novelty in such pipelines is prescribed in advance, and anomalies that fall outside those features go undetected. A human expert examining the data can attend to features beyond the predefined set and flag anomalies, which is why even systematic surveys end in expert curation (7, 13, 14). Such curation does not scale with the data and remains a bottleneck in mining these resources.

Large language models (LLMs) can now write and run code, operate analysis tools and databases, and complete long tasks with little supervision (15–18). LLMs are automating parts of software engineering and data analysis (19, 20), and accelerating biological research, where multi-agent systems have originated drug candidates, protein designs, gene-editing experiments, and mechanistic hypotheses that were confirmed at the bench (21–25). More recently, autonomous agents have planned and executed chemical syntheses, machine-learning studies, and biological data analyses end to end, from a stated objective to a written report (26–30). We reasoned that genome mining, an iterative process of collecting, classifying, and investigating sequences, could be automated in the same way with our latest frontier models. We further asked whether LLMs that reason over the data might also notice anomalies beyond the features a pipeline prescribes, supplementing manual curation.

As a test case we chose reverse transcriptases (RTs), a widespread enzyme family that canonically copies RNA into DNA, an activity that underpins biotechnologies from cDNA synthesis to genome engineering (31–34). In prokaryotes, RTs are often co-expressed with a partner protein and non-coding RNAs (ncRNAs) encoded in the same locus (35, 36). In retrons, for example, the RT copies a dedicated ncRNA into single-stranded DNA that functions with an effector protein in antiphage defense (37, 38). These effectors are diverse, including an N-glycosidase that hydrolyzes NAD^+^ in Retron-Eco1 and a membrane protein predicted to disrupt the membrane in Retron-Eco6 (37, 39, 40). New RT systems that act with novel ncRNAs and partner genes continue to be reported (36, 41–47), suggesting that more remain among RT loci that have yet to be examined.

Here, we present an autonomous genome-mining campaign in which Claude agents surveyed RTs across 1.9 billion metagenomic protein clusters. The campaign was structured so that agents could pursue their own observations beyond a predefined pipeline. One such observation led to the discovery of the array-associated RT (ART) family, when an agent reading the DNA beside a phage RT noticed an unannotated array of tandem repeats. ART belongs to no described RT class and is typified by its non-coding DNA (ncDNA) repeats reminiscent of CRISPR arrays, an architecture that may serve to diversify the function of a single enzyme. This work demonstrates that AI systems can notice anomalies in primary biological data and surface them as a human expert would, easing the curation bottleneck in genome mining at a large scale.

## Results

### Agentic genome mining with Claude

We built an agentic harness, software that coordinates LLM agents, to carry out a genome-mining campaign based on a brief that outlines research goals (**Fig. 1A**). The research brief is first converted into a series of analysis stages that represent broad steps, such as surveying a database for homologs. Each stage is carried out as smaller tasks, such as a sequence profile search, in which one Claude Code agent acting as the worker proposes a plan and executes it while a second agent acts as the supervisor to review the plan and results. The supervisor can open new tasks for new workers in response to observations made during the work, allowing the campaign to extend itself as findings accumulate. All plans, results, and reviews are written to a shared record, so that the harness operates as a multi-agent system with a common memory. The campaign ends when the task queue is exhausted, and its findings are delivered to human reviewers as written reports.

**Figure 1.**
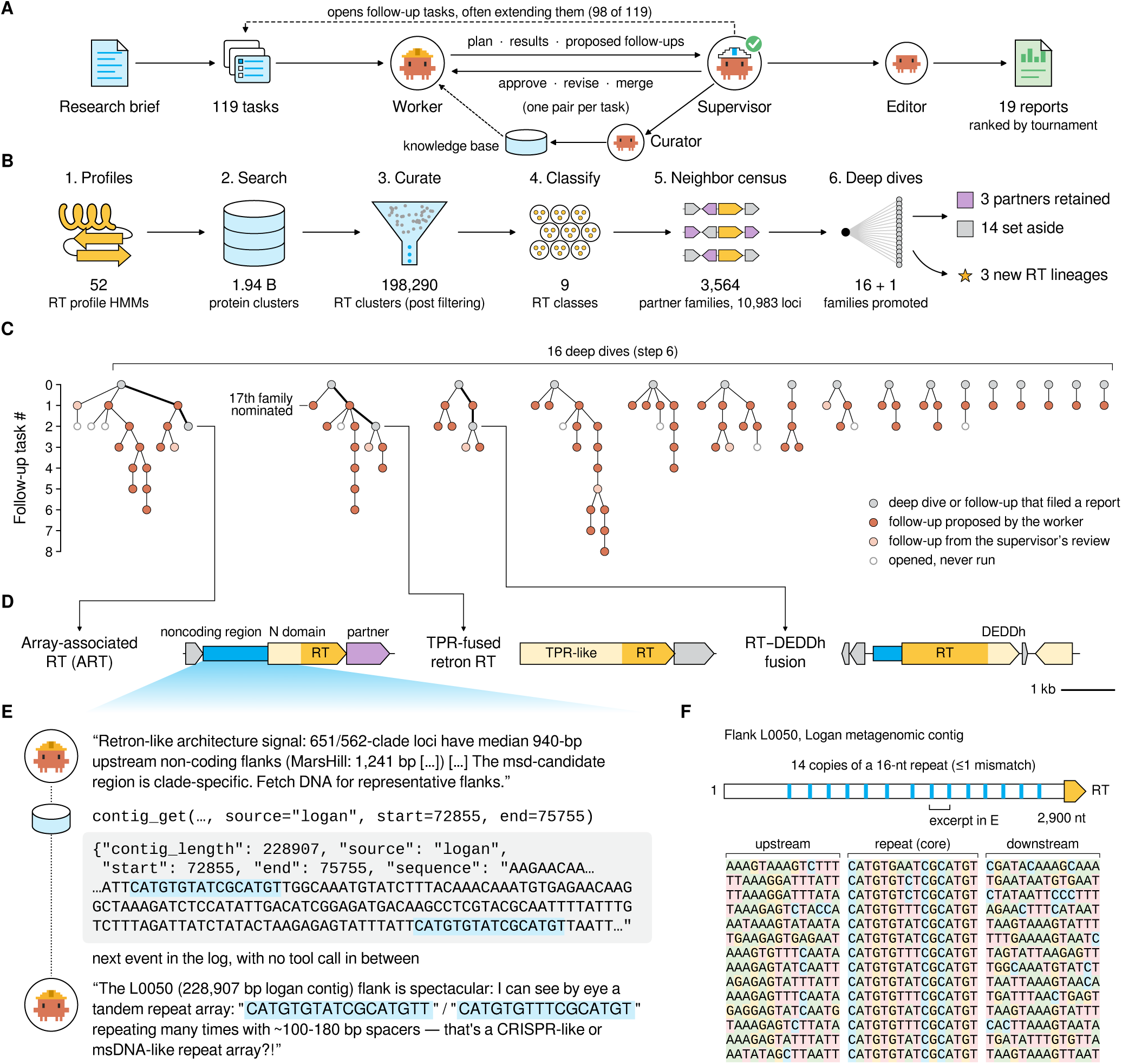
Agentic discovery of novel RT lineages. (**A**) Schematic of the harness. Instances of Claude act as workers to plan and run tasks, supervisors to review them and open follow-up tasks (98 of 119 tasks), curators to record findings in a shared knowledge base, and editors to review reports. (**B**) Pipeline for survey of RTs and their partner proteins. (**C**) Task trees for each of 16 deep dives on RT families nominated in (B). The 17th family was promoted by a follow-up task. Bold paths lead to the first report on each new RT lineage. (**D**) Representative loci of three newly described RT lineages, to scale. (**E**) Worker agent transcripts showing loading and interpretation of DNA in context identifying a repeat motif. Blue, repeat copies. (**F**) Schematic of locus examined by worker in (E), with 14 repeats and surrounding context.

We wrote a research brief that asked the agents to identify novel RT systems on the basis of new partner-gene associations (Supplementary Note 1). Running with Claude Mythos 5, the agents assembled their own sequence profile hidden Markov models (HMMs) and reference sets (Methods), recovered ∼200,000 RT clusters, and classified them into nine distinct RT classes (**Fig. 1B**). They then sampled ∼11,000 RT loci spanning every class and scored the 3,564 protein families that recurred in their neighborhoods as candidate partners. Sixteen families passed the agent-defined selection criteria and a review of their annotations, and were each investigated by a dedicated worker (**Fig. 1B**). The agents opened 98 new tasks from observations made along the way, one of which led to the promotion of a seventeenth candidate (**Fig. 1C**). The full campaign comprised 119 tasks and 949 agent sessions, amounting to 77 agent-hours and 215.6 million tokens over 21.5 hours of wall-clock time, without human intervention (**Supp. Fig. 1A**).

Of the 17 candidate partner families, only three were confirmed as previously unreported RT associations. The other 14 were rejected or set aside as annotation artifacts, parts of previously described systems, or general neighborhood residents rather than dedicated partners (**Supp. Fig. 1, B and C**). Beyond the partner-gene associations the census scored, workers also flagged unusual features of individual RTs and their flanking DNA for follow-up by new workers. Three of the flagged RTs were identified as new RT lineages (**Fig. 1D**; **Supp. Fig. 1C**). In total the campaign concluded with 19 reports, comprising 16 for the 17 candidate families (two were described in a single report) and three for the new RT lineages.

### An unexpected observation led agents to array-associated RTs

We turned our focus to the reports that ranked highly according to a tournament-style evaluation of the 19 reports (Methods). One report described a new lineage of RTs, largely confined to jumbo-phage genomes, whose defining feature lay in the ncDNA upstream of the RT (**Fig. 1D**). Since the mission was directed at partner protein-coding genes rather than ncDNA features, we examined the session transcripts to determine how the agents had identified this RT family (**Supp. Fig. 2A**). The agents initially selected this candidate because it co-occurred with a nearby phage gene, the β-like subunit of the jumbo-phage non-virion RNA polymerase. A worker agent later rejected this gene as a spurious association, but chose to queue a follow-up task to study the RT more closely. From the reasoning trace:

**Figure 2.**
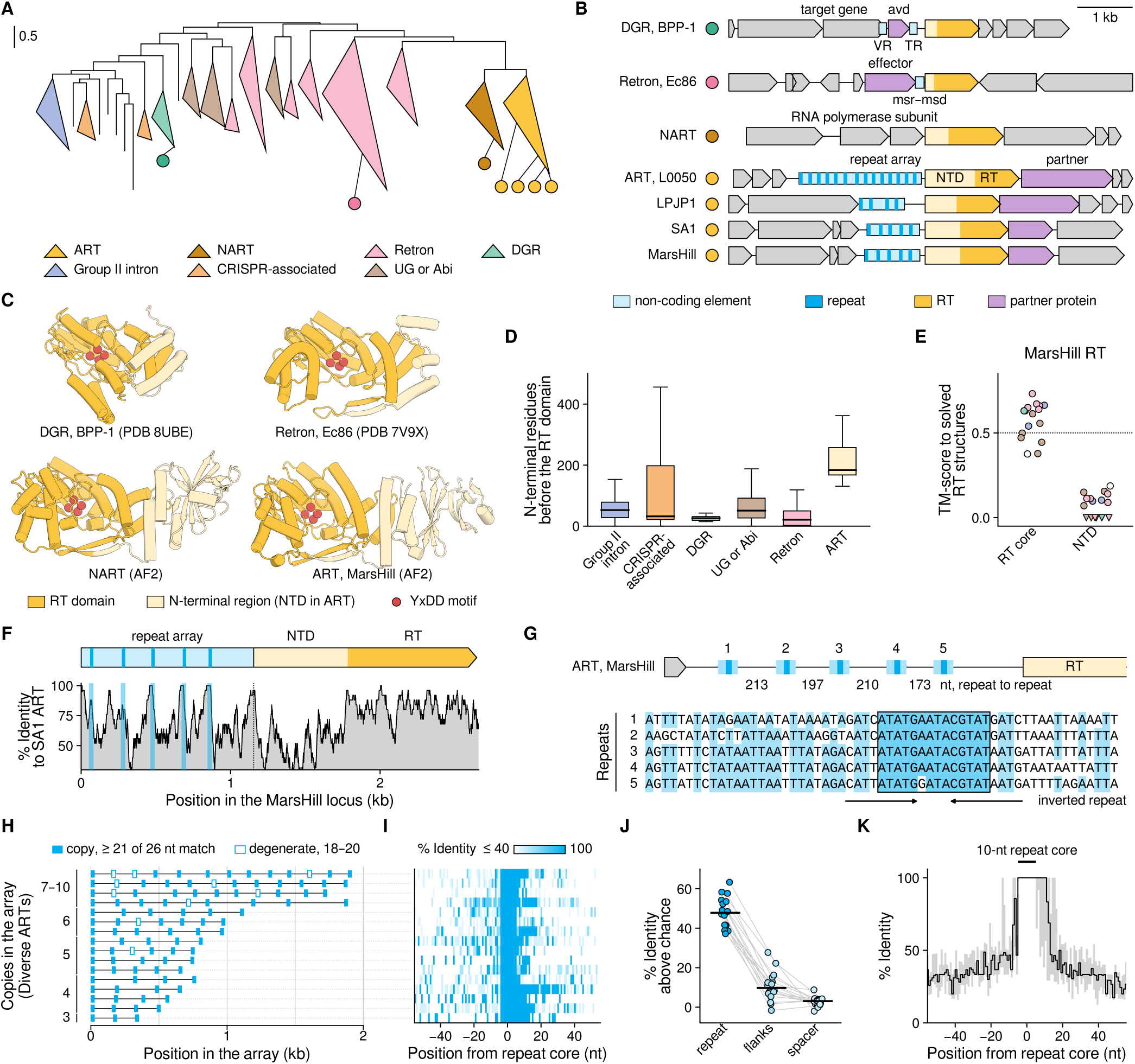
Features of array-associated reverse transcriptase loci. (**A**) Maximum-likelihood tree of 774 RT domains, midpoint-rooted and collapsed by class. (**B**) Representative loci of the RT families in (A), to scale. NTD, N-terminal domain. (**C**) Solved and predicted (AlphaFold2) structures of related RTs. (**D**) Number of N-terminal residues ahead of the RT domain of representative RT systems. Boxes, quartiles; whiskers, 1.5 times the interquartile range. (**E**) Foldseek TM-scores of the MarsHill RT core and NTD against solved RT structures (n = 16; triangles, no alignment reported). (**F**) Sequence identity between the SA1 and MarsHill ART loci (25-nt sliding window). Blue, repeats. (**G**) Repeats of the MarsHill ART, shown with flanking conserved regions. (**H**) Pattern of repeat arrays across 16 representative ART loci. Degenerate repeats are shown. (**I**) Identity between array units in (H), centered on conserved repeat core. (**J**) Sequence identity of repeats, repeat flanks, and spacers within the same array, above that of shuffled sequences. Lines join one array; bars, medians. (**K**) Sequence identity around repeat core across ART systems (n = 16 arrays; line, median; band, interquartile range).

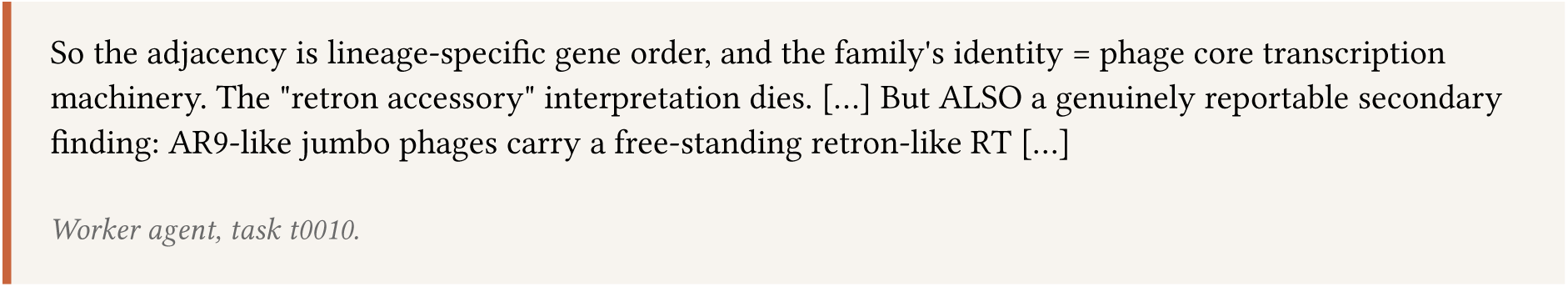

### Worker agent, task t0010

Retrons typically act in phage defense, so the agent considered the presence of the retron in the jumbo-phage genome to be noteworthy. Because retron RTs act with an ncRNA encoded upstream, the agent’s supervisor asked the next worker to check the 5′ flanks of these RTs for such an RNA. The RTs beside the polymerase gene had only short non-coding regions upstream (median 27 bp), too short to encode an RNA, whereas their closest relatives had long ones (median 940 bp). The worker read the upstream sequence of these relatives directly into its context, which led to the following reasoning trace (**Fig. 1E**; **Supp. Fig. 2B**):

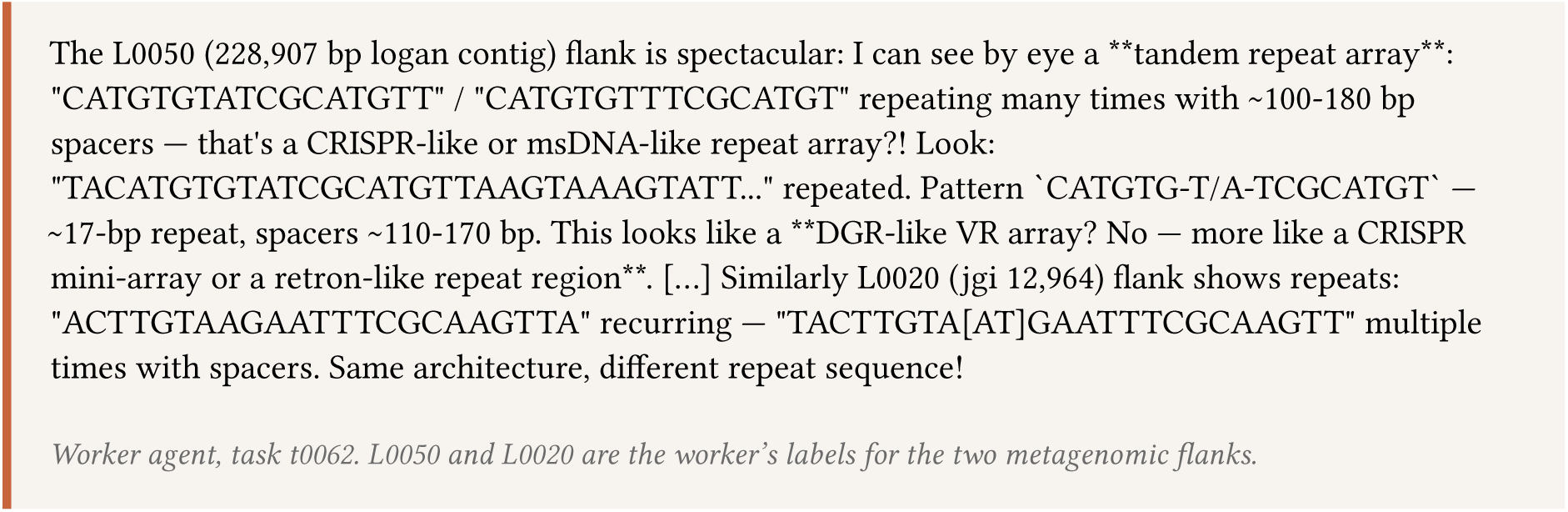

### Worker agent, task t0062. L0050 and L0020 are the worker’s labels for the two metagenomic flanks

The worker directly read this tandem repeat array pattern as a potentially undescribed RT feature. The subsequent reasoning trace shows evidence that the worker then questioned the novelty of this system, comparing it with known RT systems:

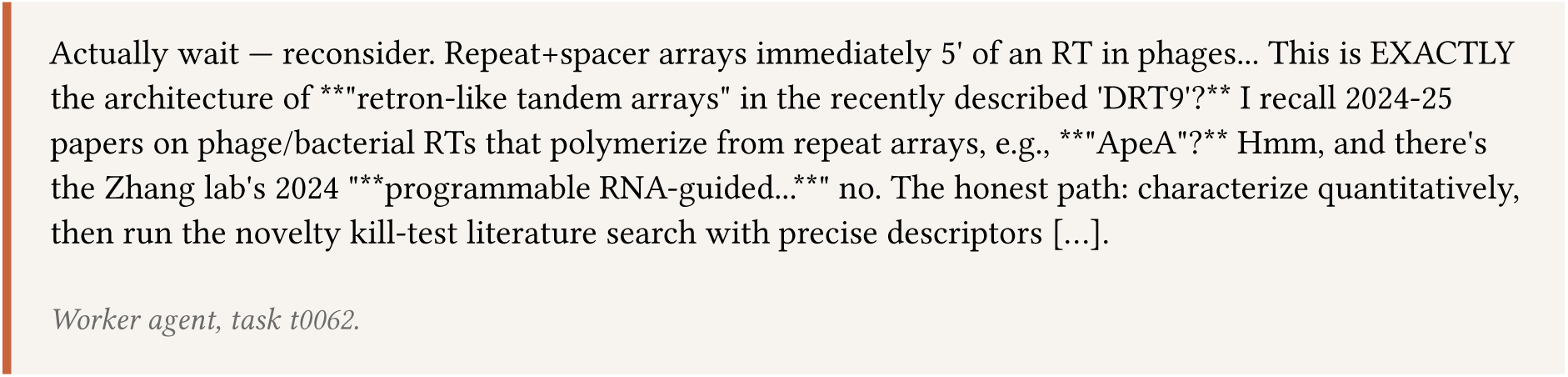

### Worker agent, task t0062

The worker subsequently analyzed the non-coding regions and confirmed they were different from any known non-coding elements, including CRISPR arrays (**Supp. Fig. 2C**). A repeat-counting script written by the worker found one locus with 14 copies of a 16-nt repeat in the region, separated by unique spacers of 100 to 200 nt (**Fig. 1F**). A literature search later confirmed that the array did not fit any literature reported feature, and the agents filed a report for human review. We named this lineage of jumbo-phage RTs array-associated RTs (ART) after the array that defines it.

### Array-associated reverse transcriptases (ART) are a new family of RTs

We next determined in interactive Claude Science sessions whether the array and its RT define a coherent family beyond the loci the agents had examined (Methods). Profile HMM searches of our metagenomic database and public genomes identified 95 distinct RT clusters (90% identity) in cultured jumbo phages and predicted viral contigs, 28 of which carried a detectable repeat array upstream of the RT (**Supp. Fig. 3A**; Methods). Phylogenetic analysis placed the RTs in a clade next to retrons, in line with the agents’ initial classification (**Fig. 2A**; **Supp. Fig. 3B**). In these loci, the array lies upstream of the RT, a feature missing from other RT systems, and a dedicated partner gene lies directly downstream, both of which the agents described in their report (**Fig. 2B**; **Supp. Fig. 3C**).

**Figure 3.**
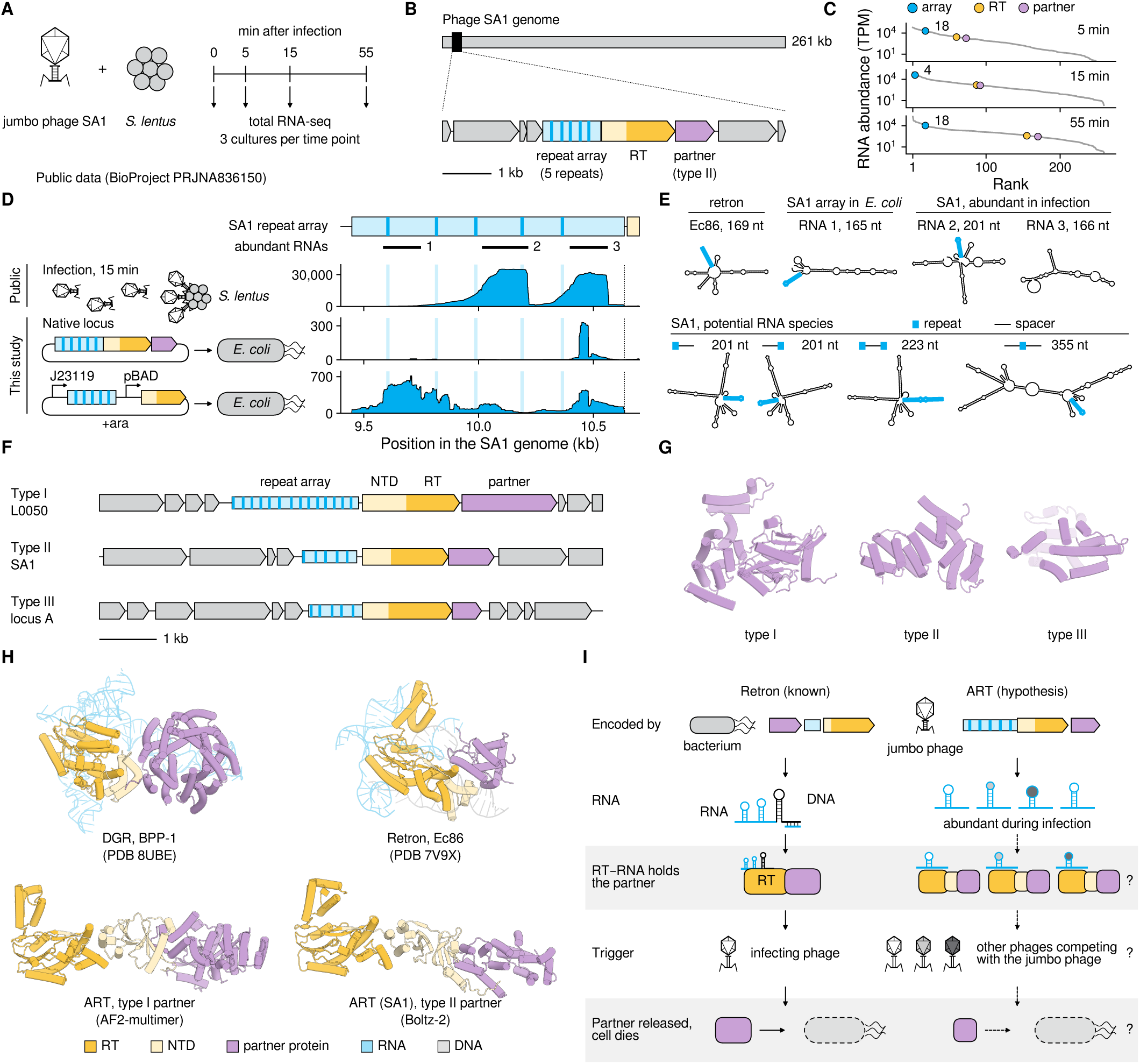
ART arrays are transcribed into abundant RNAs, and ART family RTs co-occur with dedicated partner proteins. (**A**) Schematic of the SA1 phage infection time-course experiment in *S. lentus*, performed with RNA-seq for BioProject PRJNA836150. (**B**) Schematic of phage SA1 genome and resident ART locus. (**C**) SA1 transcript abundance during infection. TPM, transcripts per million. Numbers, rank of the array RNA. (**D**) Alignments of RNA-seq traces to SA1 ART array. (**E**) Secondary structures of Ec86 retron RNA compared to putative SA1 RNA repeats. (**F**) Representative ART loci with divergent partner proteins. (**G**) Predicted structures of the three types of observed ART partner proteins (59, 7, and 3 lineages; AlphaFold2-multimer for type I, Boltz-2 for types II and III). (**H**) Structures of RT-partner protein interactions compared to putative co-folds of ART family RT and partners. (**I**) Putative model of ART biological function compared to characterized retron function. Dashed arrows, hypothesized steps.

The RT itself has an unusually long N-terminus, with about 180 residues preceding the polymerase domain where other RTs typically carry about 50 or fewer (**Fig. 2, C and D**). The polymerase domain retains the catalytic YxDD motif in all 93 members whose sequences span it (two are truncated before it) and superimposes on solved retron and diversity-generating retroelement (DGR) RT structures (**Supp. Fig. 4, A to C**). In contrast, the N-terminal extension of the RTs shows at most weak similarity to any known sequence family or fold, and is the most variable region of the protein among members (**Fig. 2, E and F**; **Supp. Fig. 5A**). Nevertheless, it is present in every member of the family and constitutes the second hallmark of ART RTs (**Supp. Fig. 5B**). The polymerase-adjacent RTs that first led the agents to ART form its sister clade (**Fig. 2, A and B**). These RTs also have a long N-terminus (**Supp. Fig. 5C**), yet none of their 17 assessable loci carries a repeat array and none encodes an ART partner (**Supp. Fig. 5, D and E**). We therefore refer to them as non-array-associated RTs (NART). Their N-terminus adopts a single predicted fold distinct from those of ART (**Supp. Fig. 5, F to H**).

**Figure 4.**
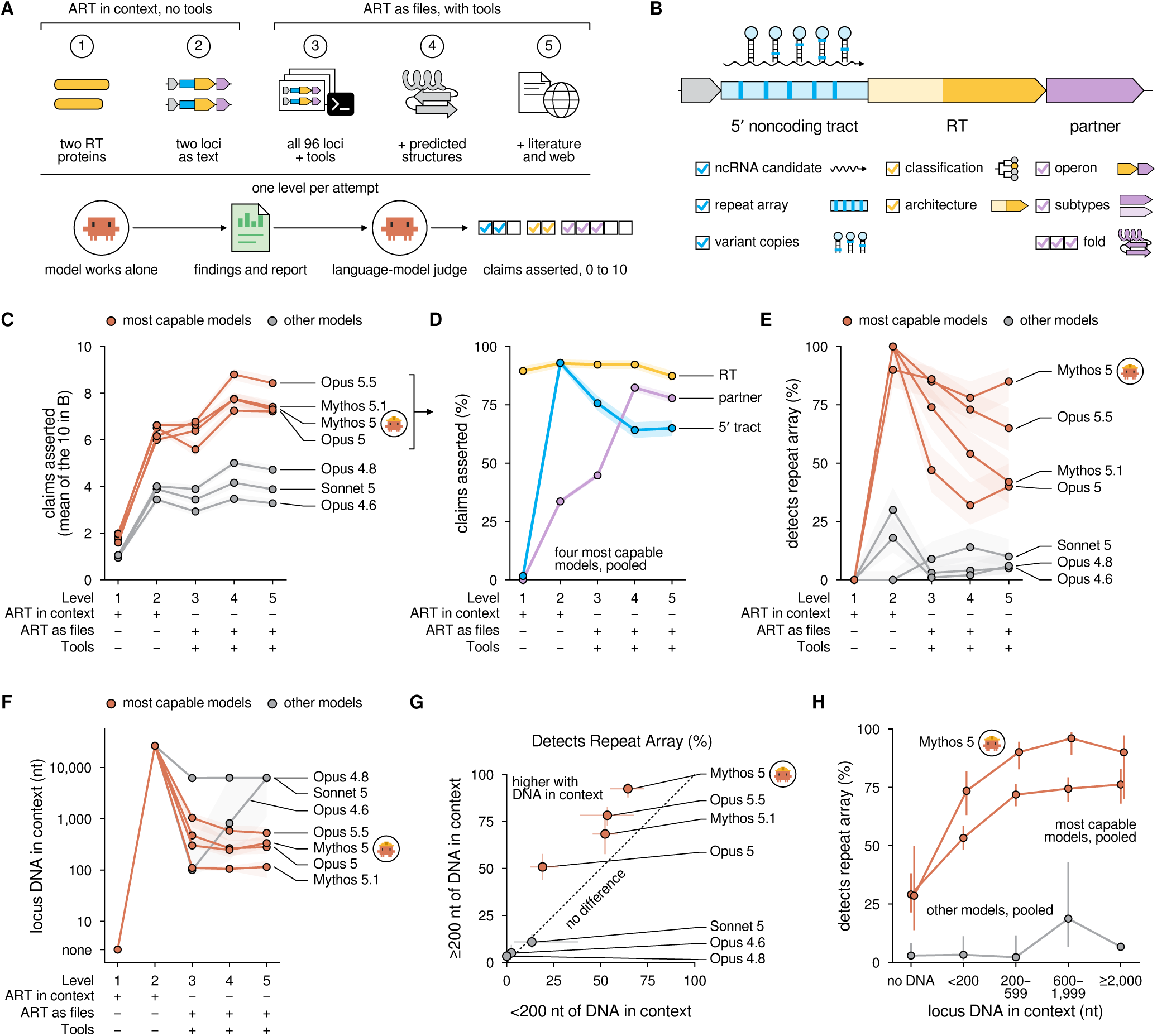
The most capable models recognize the ART array, and recognition depends on reading DNA into context. (**A**) The fixed-input benchmark. Each attempt receives one of five levels of information, with ART sequences in the model’s context and no tools (levels 1 and 2) or as files with analysis tools (levels 3 to 5). The model works alone, and a language-model judge scores its findings and report on ten claims. (**B**) The ten claims, grouped by the part of the locus they concern. Each box is one claim. (**C**) Mean number of rubric claims asserted at each level by seven models (n = 100 attempts per model and level). Orange, the four most capable models. Gray, the other three. Icon, the model of the original campaign. Shading, 95% confidence intervals. (**D**) Claims asserted about each part of the locus as a percentage of the total number of claims, for the four most capable models pooled. (**E**) Attempts that recognized the repeat array. (**F**) Median longest contiguous stretch of locus DNA in the model’s context. (**G**) Recognition of the array at levels 3 to 5 in attempts with at least 200 nt of contiguous locus DNA in context, against attempts with less. Dashed line, no difference. (**H**) Attempts at levels 3 to 5 that reported the repeat array, by the longest contiguous stretch of locus DNA in the model’s context, for the four most capable models pooled (n = 110, 360, 338, 266, and 126 attempts per bin) and the other three pooled (n = 103, 61, 45, 16, and 675). Bars, 95% confidence intervals.

**Figure 5.**
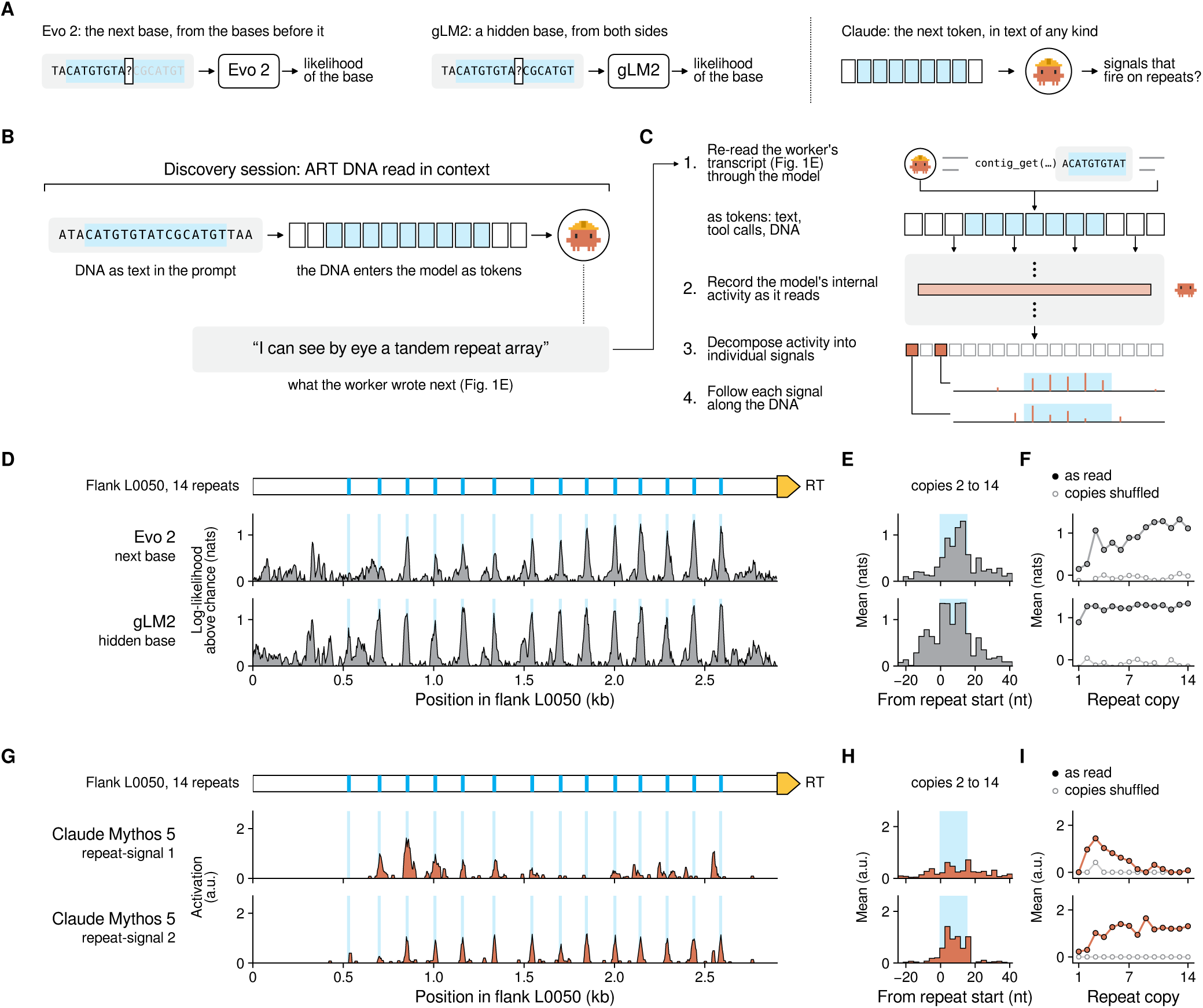
Internal Mythos 5 signals respond to repeats as the sequence is read. (**A**) How each model reads DNA. Evo 2 predicts the next base, gLM2 a hidden base, and Claude the next token of any text. (**B**) In the discovery session the ART DNA was in the prompt and entered Claude as tokens. Quote, what the worker wrote next (Fig. 1E). (**C**) Readout of internal signals. The worker transcript of Fig. 1E was passed through Mythos 5 again, and its activity at one layer was decomposed into individual signals. (**D**) Genomic models along flank L0050 (Fig. 1F). Log-likelihood of the true base above chance (16-nt running mean). Evo 2 was scored through Evo Designer. (**E**) Aligned on the repeat, mean of copies 2 to 14 in 3-nt bins. (**F**) Mean per copy, as read and with each copy shuffled in place. (**G to I**) As D to F, for two of the 12 Mythos 5 signals preselected on synthetic repeats, repeat-signal 1 and repeat-signal 2 (activation; G, 16-nt running mean). Neither is specific to DNA.

We next investigated the non-coding arrays that typify ART systems. The 28 arrays adjacent to an RT span 0.3 to 4.1 kb and consist of 3 to 21 copies of a short repeat at near-constant spacing (**Fig. 2, G and H**). Repeats are 15 to 49 nt long, each with a palindromic core of ∼15 nt and less conserved edges, and are shared within but not across ART clades (**Fig. 2G**). The spacers between repeats are 120 to 220 nt long and unrelated to one another within an array (**Fig. 2, H to K**). Between related phages such as MarsHill and SA1, spacers are retained in order and diverge at a similar rate to the RT gene (**Fig. 2F**). Across ARTs, conservation of repeats is generally higher than conservation of spacer sequences. These features contrast with CRISPR arrays, where near-identical repeats with sharp boundaries alternate with spacers of only ∼30 nt that are gained and lost between related strains. Critically, no *cas* genes occur near any ART locus, suggesting that the arrays represent a new type of non-coding repeat element.

### ART repeat arrays are highly expressed during phage infection

To learn more about the array, we prompted a Claude Science session to search for further information on ART loci. The session autonomously identified and retrieved RNA reads from a published infection time-course study of *Staphylococcus* phage SA1 (48), which contains an ART system (**Fig. 3, A and B**). This dataset revealed that the ART repeat arrays are highly expressed at early, mid, and late stages of infection (**Fig. 3C**; **Supp. Fig. 6, A and B**). Remarkably, at 15 minutes post infection, the array-derived RNAs account for up to 8% of phage RNAs, making them among the most abundant phage transcripts (**Fig. 3C**). Within the array, the transcripts resolve into shorter species whose boundaries are reproducible across replicates (**Fig. 3D; Supp. Fig. 6A**).

To examine the array RNA independently of phage infection, we expressed the SA1 ART system on plasmids in *Escherichia coli* and performed small-RNA sequencing. Across both a native locus and a heterologous-promoter construct, we observed discrete short RNAs from the array similar to those formed during infection (**Fig. 3D**). Folding predictions for array units placed the repeat in a base-paired stem regardless of how each unit was cut from the array, consistent with the repeat itself containing a short inverted repeat (e.g., a palindrome) (**Fig. 2G**; **Fig. 3E**; **Supp. Fig. 6C**). These results suggest that a single ART array gives rise to several distinct short ncRNAs with the potential to form secondary structures.

These RNAs raised the possibility that ART is a retron-like system that operates with a bank of distinct RNAs. This prompted us to investigate the ART partner genes, since the partner protein executes the system’s function in retrons. We found three unrelated partner families that each occur nearly exclusively beside ART family RTs, and classified ART systems accordingly (**Fig. 3, F and G**; **Supp. Fig. 7A**). Type I, found at 59 of 95 loci across the *Listeria* phage and environmental lineages, encodes a protein of about 600 residues with two tandem GCN5-related N-acetyltransferase (GNAT)-like folds (**Supp. Fig. 7, B to D**). Type II, confined to the *Staphylococcus* phage clade, encodes an all-helical protein of about 270 residues, and type III encodes a smaller helical protein of about 170 residues in one environmental sub-lineage; neither appears to have clear homologs in protein domain or structural databases (**Supp. Fig. 7, B and D**). The three partner families share no detectable similarity in sequence or predicted structure, suggesting that ART family RTs have been paired with unrelated partners more than once. Structure predictions of cognate type I and II pairs converged with moderate confidence (interface predicted template modeling [ipTM] score ∼0.6) on an interface between the N-terminal extension of the RT and the partner (**Fig. 3H**). The N-terminal extension adopts different predicted folds in the two lineages, raising the possibility that it acts as a partner-binding module that co-evolves with its partner (**Supp. Fig. 5, F to H**; **Supp. Fig. 7, E and F**).

Taken together, ART loci share the three components of a retron (an ncRNA, an RT, and a dedicated partner) but carry a bank of distinct RNAs where retrons carry one (**Fig. 3I**). In retrons the RT and the single-stranded DNA that it produces (msDNA) hold the effector in an inactive complex until phage infection or damage to the msDNA releases it, killing the cell (37, 38). We hypothesize that ART may work on a similar principle, with each unit RNA assembling its own complex with the RT and partner, so that one enzyme and effector pair yields several complexes that differ only in their RNA (**Fig. 3I**). The differences between these RNAs may in turn allow the system to respond to different triggers, given that phage proteins trigger retrons by directly perturbing the msDNA (38).

### The most capable models recognize the ART array when they read its DNA

We next asked whether the discovery of ART was reproducible within our harness, so we ran the same campaign ten more times. Nearly every campaign that completed the census sampled ART loci, and in two campaigns, workers investigated the lineage as follow-up. However, none read the DNA upstream of the RTs, and the array was missed in every rerun. We attribute these results to the broad protein search space of RTs and the non-deterministic behavior of the harness.

We therefore built a set of fixed-input benchmarks in which a model is given ART sequences directly in context or in a filesystem with tools, and scored on its ability to analyze and accurately describe the system (**Fig. 4A**; Methods). Every model was tasked with characterizing the ART system from the input provided and reporting what it deemed important. The resulting report was scored by a judge model (Mythos 5) against ten curated features that we deemed characteristic of the ART family (**Fig. 4B**). Testing seven Claude models, we found a clear gap between the four most capable models (Opus 5.5, Mythos 5.1, Mythos 5, and Opus 5) and the other three (Opus 4.6, Opus 4.8, and Sonnet 5) (**Fig. 4C**). This suggests that the original search campaign, run on Mythos 5, benefited from these improved capabilities.

We then broke down our benchmarks to look at how the most capable models performed at characterizing the three main components of ART: the array, the RT, and the partner (**Fig. 4, D and E**). We noticed that the RT was accurately described by the most capable models, even with just the protein sequences of two RTs provided. Comparatively, more tools and information helped the models better describe the partner gene. Surprisingly, more information and tooling inhibited the models’ ability to accurately describe the array, as was accomplished in the original search campaign. Given only the loci in context, the most capable models accurately described the array in at least 90% of attempts, compared to as low as 32% (Opus 5, level 4) with tooling.

We hypothesized that this lower performance stems from these models failing to read the attached sequences into their context, so we inspected transcripts for the amount of DNA each attempt read (**Fig. 4F**). With files, 39% of their attempts never read a contiguous stretch of 200 nt or more and therefore never saw more than about one repeat unit. When a stretch of at least 200 nt was read, recognition was 16 to 32 percentage points higher for each of these models (**Fig. 4G**). As more DNA sequence was read into the context of the model, recognition of the array rose from 29% to as high as 76% of attempts for the four models pooled, and as high as 96% for Mythos 5 (**Fig. 4H**). These results indicate that direct observation of the DNA by the model is critical for recognizing the array, as it was in the original campaign.

### Mythos 5 internal signals respond to DNA repeats

Genomic language models have been shown to learn representations of repeated DNA (**Fig. 5A**). These models can be used to detect repeat motifs in DNA (10), their embeddings can distinguish repeat families (49, 50), and a signal that responds to the spacers of CRISPR repeat arrays has been isolated in Evo 2 (49). We wanted to determine if Claude models have similarly learned to represent and recognize DNA repeats like those found in ART arrays (**Fig. 5B**). To ask how the model’s internal activity responds as it reads DNA, we used the transcript of the original discovery session to decompose that activity into interpretable signals (**Fig. 1, A and E; Fig. 5C**).

Since genomic language models can detect repeats, we ran Evo 2 and gLM2 on the same L0050 ART locus Mythos 5 explored during the initial campaign (**Fig. 1F**). The log-likelihood profiles of the genomic language models consistently identified the repeats with high confidence (**Fig. 5D**). These profiles were centered on the ART repeat and lost when the nucleotides of each copy were shuffled in place (**Fig. 5, E and F**). gLM2, which reads the sequence on both sides of a base, predicted every copy including the first. Evo 2, which reads only the preceding sequence, predicted the first two copies poorly and improved along the array.

Two Mythos 5 signals (repeat-signal 1 and repeat-signal 2) responded to the array in a similar fashion to the models trained only on biological sequences (**Fig. 5, G to I**). Both were silent before the array, and only repeat-signal 2 weakly fired at the first repeat copy. Repeat-signal 1 peaked at the third copy and faded along the array, with activity scattered across the copies and flanking sequences (**Fig. 5, G and I**). In contrast, repeat-signal 2 strengthened by the third copy and remained steady thereafter, with activity largely confined to the repeat and the nucleotide that follows it (**Fig. 5H**). Shuffling each copy in place silenced repeat-signal 1 on 12 of 14 copies and repeat-signal 2 on all 14 copies (**Fig. 5I**). These results indicate that the repeat triggered a signal inside the model as the ART array sequence was read. This was immediately followed by natural language reasoning that named the sequence a “tandem repeat array,” which enabled the initial identification of ARTs.

## Discussion

Many scientific discoveries originated when a scientist looked closely at the data and recognized something unusual as worth pursuing. Here, these steps were taken by AI agents working together without human supervision. The array that defines ART was recognized after the model read the raw DNA sequence into its context. Both our benchmarks and the Mythos 5 internal signals that respond to repeated DNA suggest that this “genomic vision” directly enabled the discovery. Paired with a harness that let agents pursue and refute their own hypotheses, this capability allowed an AI system to take the first steps of a biological discovery on its own.

These observations come at a time of rapid progress by AI systems in mathematics. This is exemplified by LLM-derived solutions to long-standing open problems (51–53). Whereas a mathematician can work entirely from first principles, a biologist often must test their ideas in the laboratory (54, 55). Much can be inferred from sequence data alone, but our findings on ART await experimental characterization. ART is a new phage-encoded member of the growing list of prokaryotic RT systems that are typified by ncDNA repeats that bear striking resemblance to CRISPR arrays. Arrays of ncRNAs were recently discovered beside the unrelated UG27 RTs using a bespoke genome language model developed to mine non-coding elements (56), suggesting that this architecture arose more than once in nature. Here, the ART array was recognized by a general-purpose model directly reading the DNA instead. Unlike CRISPR spacers, the ART units are conserved between related phages, and they are transcribed in large excess over the enzyme, suggesting one enzyme supplied with a repertoire of templates or baits (37, 38, 41, 42). Beyond these observations, we have not shown that the RT is active or that the unit RNAs are its substrates. Whether the RT and its partner interact, and what the system does for the phage, are currently unknown.

Awareness of anomaly is proposed to be at the start of scientific discovery (57), and an anomaly exists only against an expectation. For statisticians that expectation is an explicit model of the data, whereas for biologists that expectation is often derived from contextual and tacit knowledge (58, 59). In earlier attempts to automate scientific discovery, both the expectation and the judgment of novelty were supplied by human experts (54, 55, 60). Recent AI-powered methods allow agents to screen their own hypotheses for novelty, but a scientist still sets the expectation and makes the final judgment (21, 23, 24, 27–30). Here, the agents supplied both the expectation, drawn from their own knowledge rather than their task, and the judgment that the atypical observations merited further pursuit.

Agents that read primary data directly can provide the judgment that limits discovery wherever data have outgrown the attention of human scientists. The MarsHill ART locus is an example, as its original genome report identified the RT and proposed a 5′ ncRNA, yet described neither the repeats nor the partner gene (61). Our results show that agentic harness configurations and model capabilities can dramatically affect research outcomes and successful discovery. We expect that as models improve, human scientists will be enabled to explore a wider range of ideas, focus on experimental design and validation, and discover many more systems like ART by using AI systems to make connections between the vast stores of human-produced scientific knowledge.

## Supplementary Figures

**Supp. Fig. 1.**
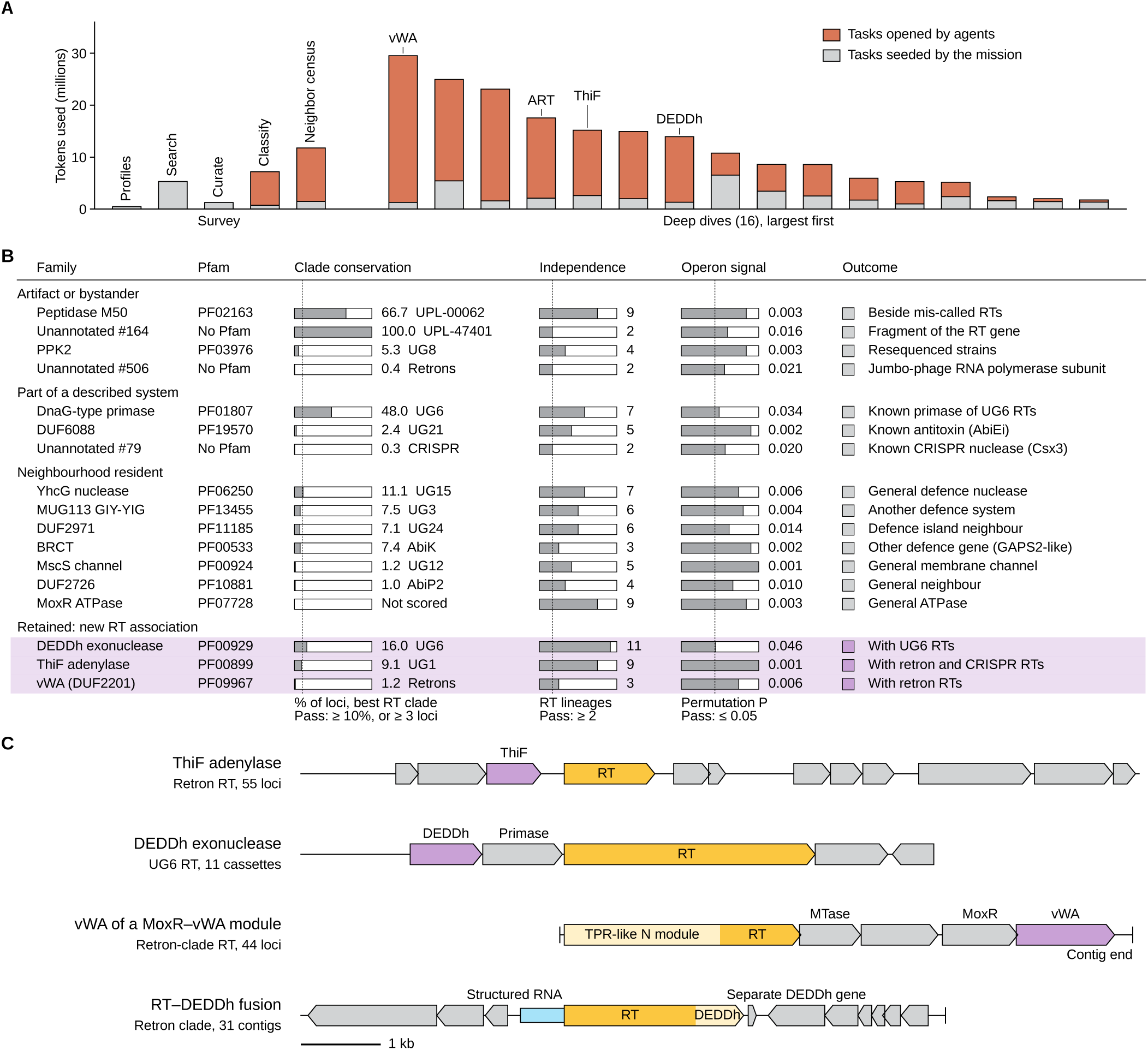
Outcomes of the promoted partner families and other new RT lineages. (**A**) Tokens used by each survey step of Fig. 1B and by each of the 16 deep dives together with all follow-up tasks that descend from it. (**B**) The 17 candidate partner families promoted from the neighbor census, grouped by final outcome, with their scores on the three promotion filters. Dotted lines, pass thresholds. (**C**) Representative loci of the three retained partner families and of the RT-DEDDh fusion.

**Supp. Fig. 2.**
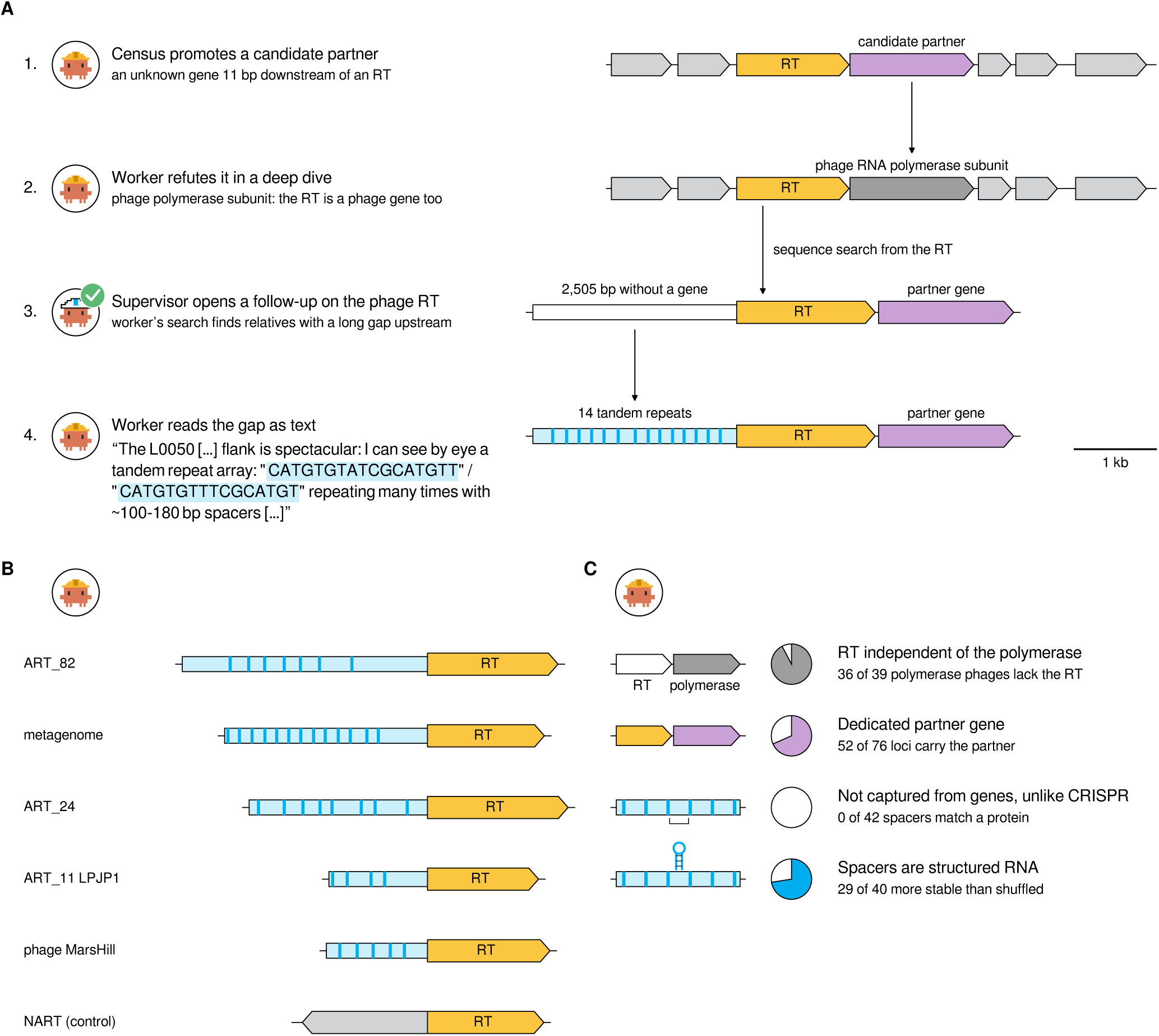
Path of the agents to the ART locus. (**A**) The four steps from a census candidate to the ART repeat array locus. (**B**) Upstream regions of further relatives retrieved by the worker, each with its own repeat array, and a control RT from the polymerase-linked lineage without one. (**C**) Tests run by the agents to refute the novelty of ARTs.

**Supp. Fig. 3.**
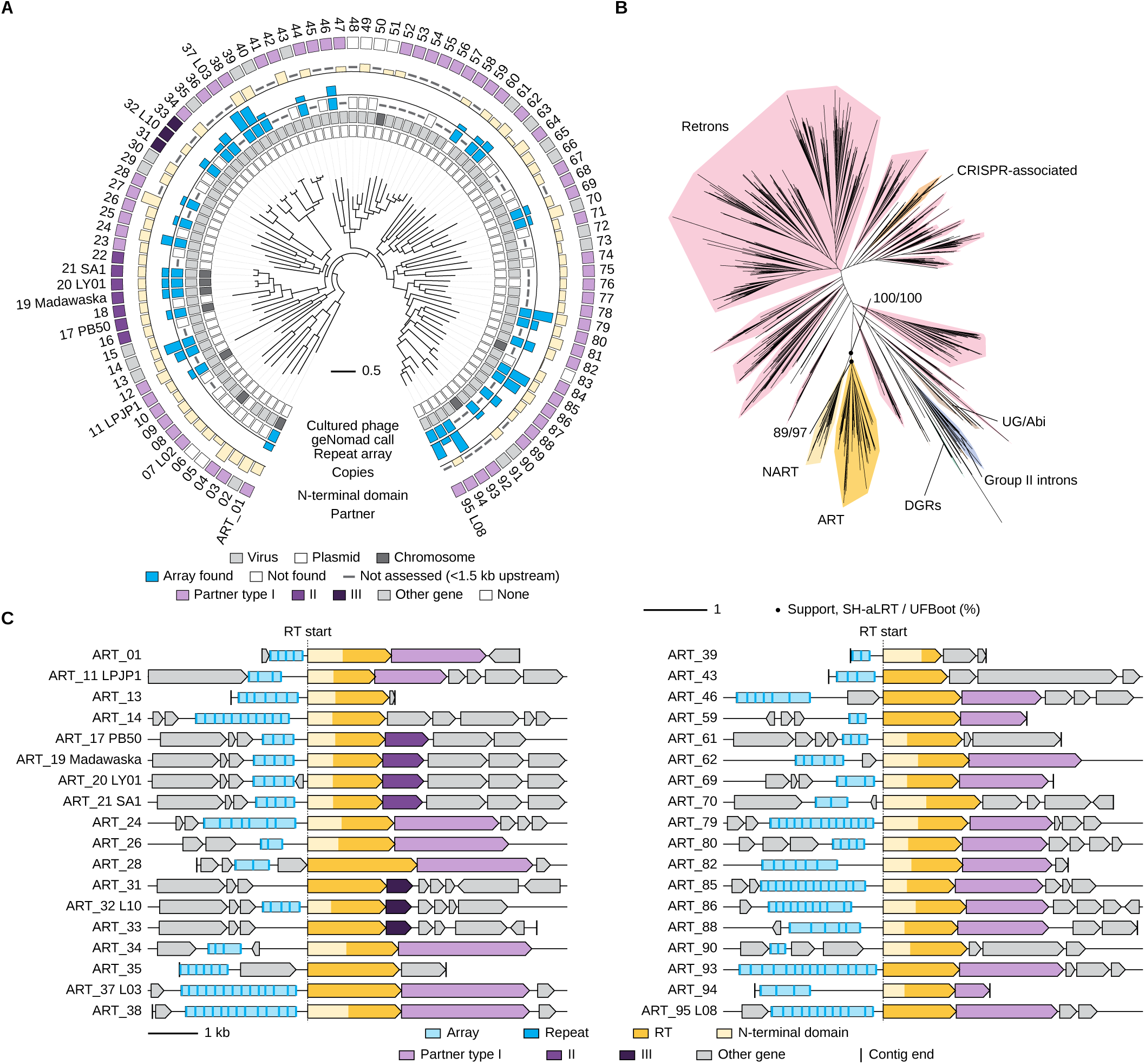
The ART family and its place among RTs. (**A**) Tree of the 95 ART family members. Dashes, not assessed (less than 1.5 kb of contig upstream of the RT) or not measured. (**B**) Unrooted maximum-likelihood tree of 774 RT domains with RT classes shaded. (**C**) Schematics of diverse ART loci.

**Supp. Fig. 4.**
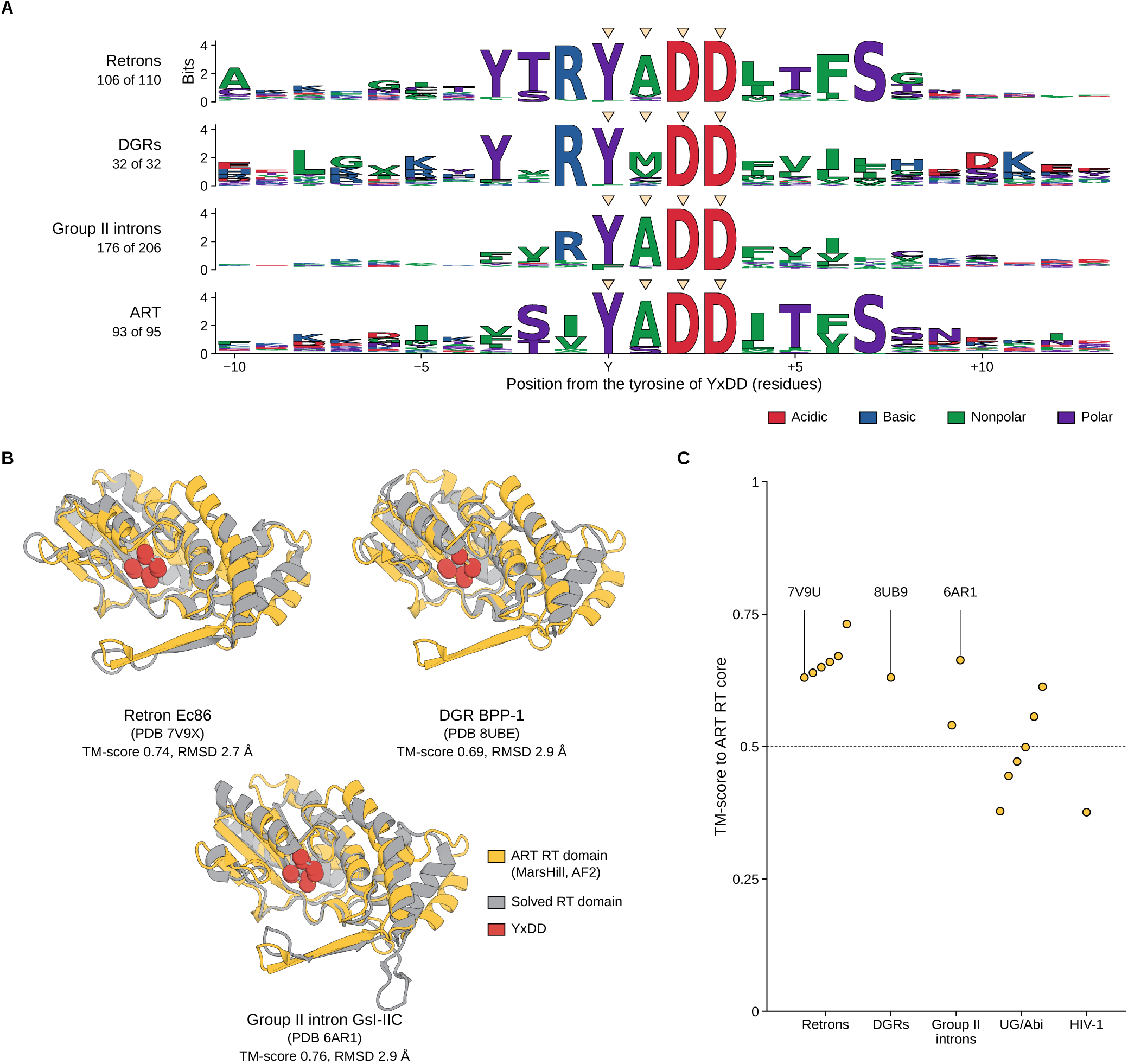
The ART family RT domain carries the catalytic motif and fold of characterized RTs. (**A**) Sequence logos of the region around the catalytic YxDD motif in retron, DGR, group II intron, and ART family RTs. (**B**) AlphaFold2 model of the MarsHill ART family RT domain (gold) superposed on solved RT domains (gray). Red spheres, Cα atoms of YxDD. (**C**) TM-scores of the ART family RT core against 16 solved RT structures, by class.

**Supp. Fig. 5.**
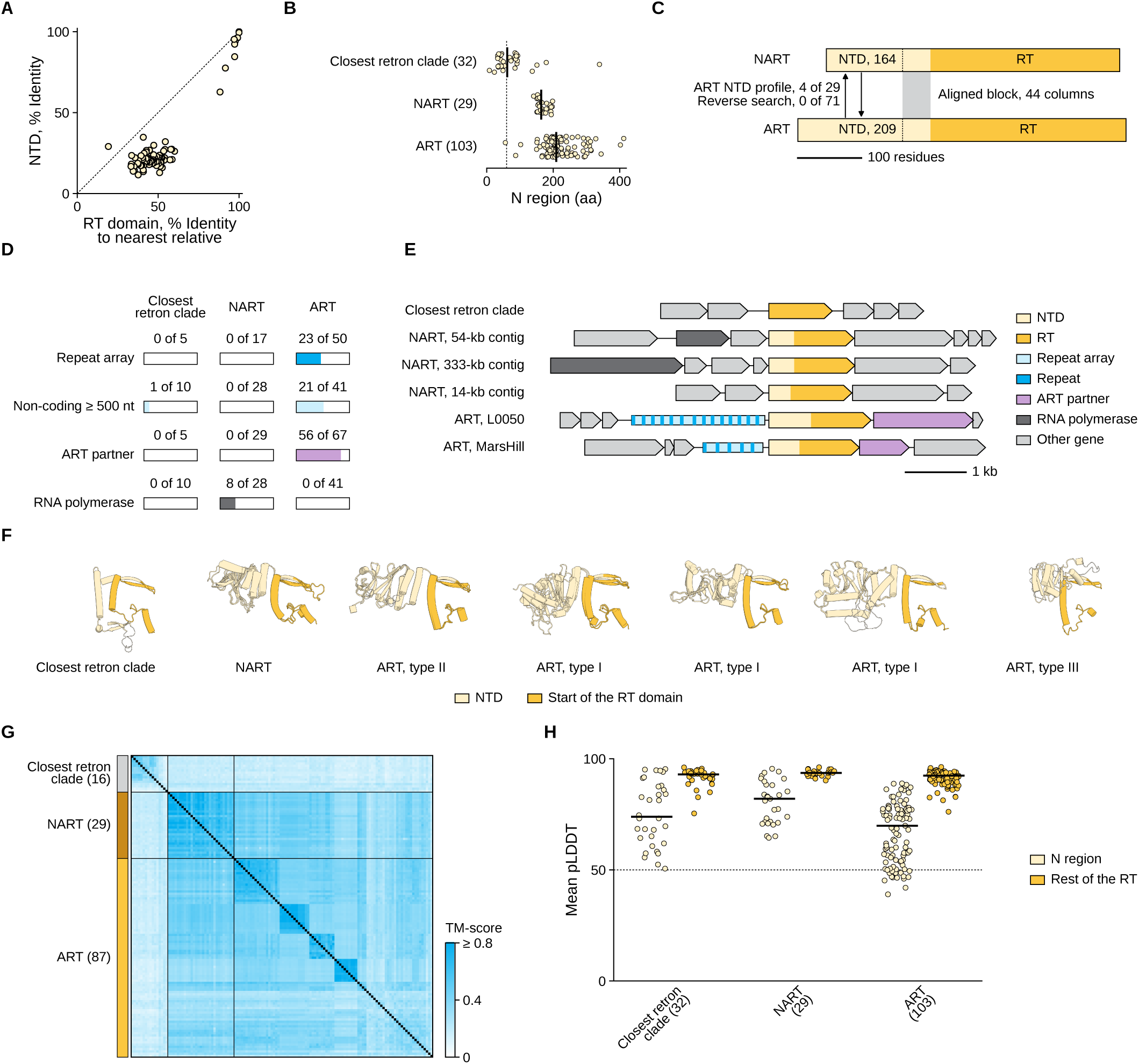
The N-terminal domain (NTD) of ART and NART family RTs. (**A**) Identity of the NTD against identity of the RT domain, both to the locus with the most similar RT domain. Dotted line, y = x. (**B**) Length of the N-terminal region of retron, ART, and NART RTs. (**C**) Comparison of median-length ART and NART RTs. (**D**) Loci with each feature among those assessable, for retron, ART, and NART RTs. (**E**) Example loci, to scale and aligned at the RT start. (**F**) AlphaFold2 models of the N-terminal region (cream) and RT-domain start (gold) for representative retron, NART, and ART RTs, with ART examples for each partner type (three structural clusters for type I). (**G**) Pairwise TM-scores between the 132 N-terminal regions of at least 60 residues with mean pLDDT of 50 or more, normalized by the longer region and ordered by group and structural cluster. (**H**) Mean pLDDT of the N-terminal region and of the rest of the RT in each model, by clade. Bars, medians; dotted line, 50.

**Supp. Fig. 6.**
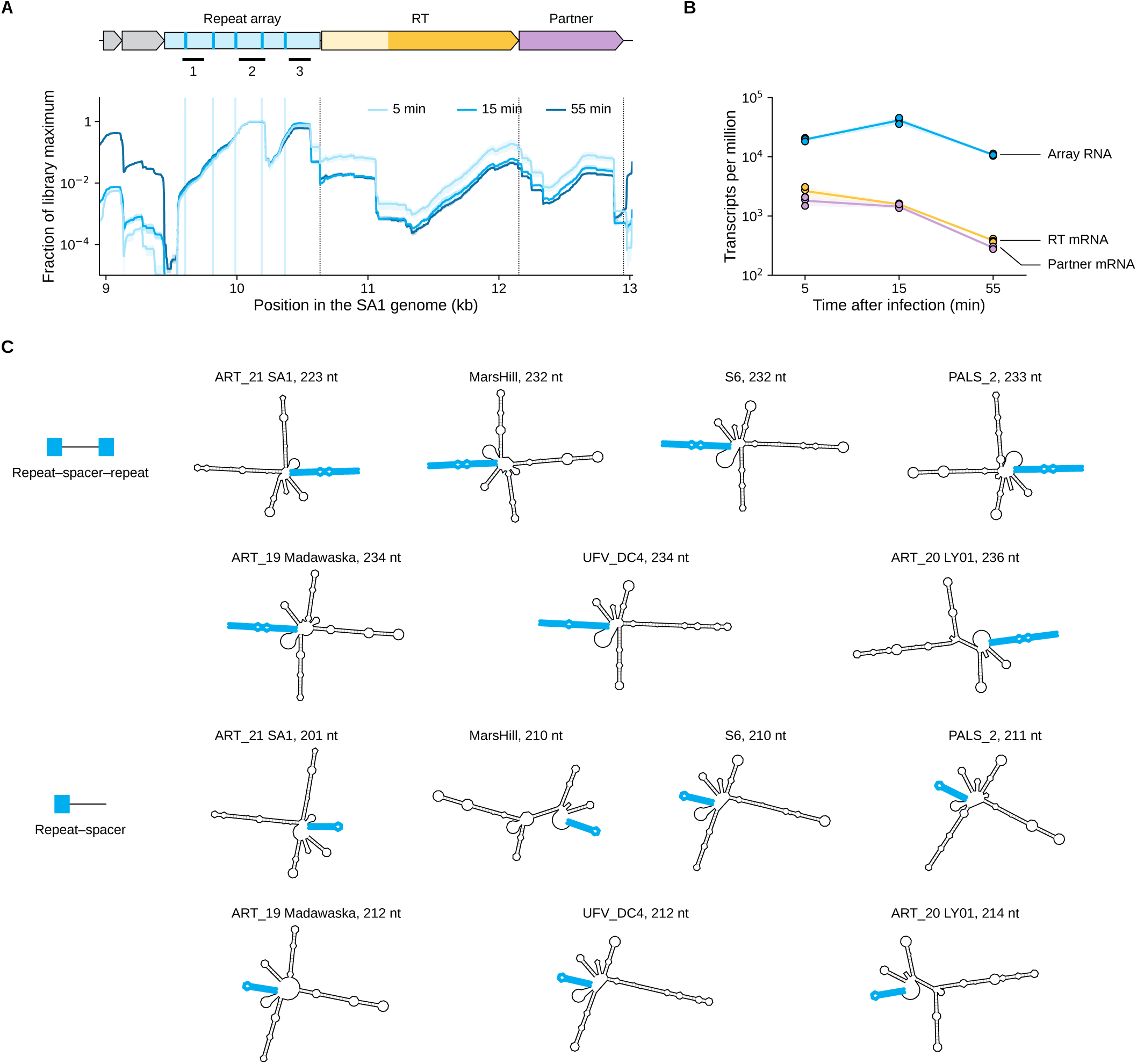
Array RNAs of phage SA1 during infection and predicted folds of ART array units. (**A**) Sense-strand RNA-seq coverage of the SA1 ART locus at 5, 15, and 55 min after infection (BioProject PRJNA836150). (**B**) Abundance of the array RNA, the RT mRNA, and the partner mRNA. TPM, transcripts per million. (**C**) Predicted minimum-free-energy structures, at one scale, of one repeat-spacer-repeat and one repeat-spacer unit from each of seven type II ART arrays. Repeats in blue.

**Supp. Fig. 7.**
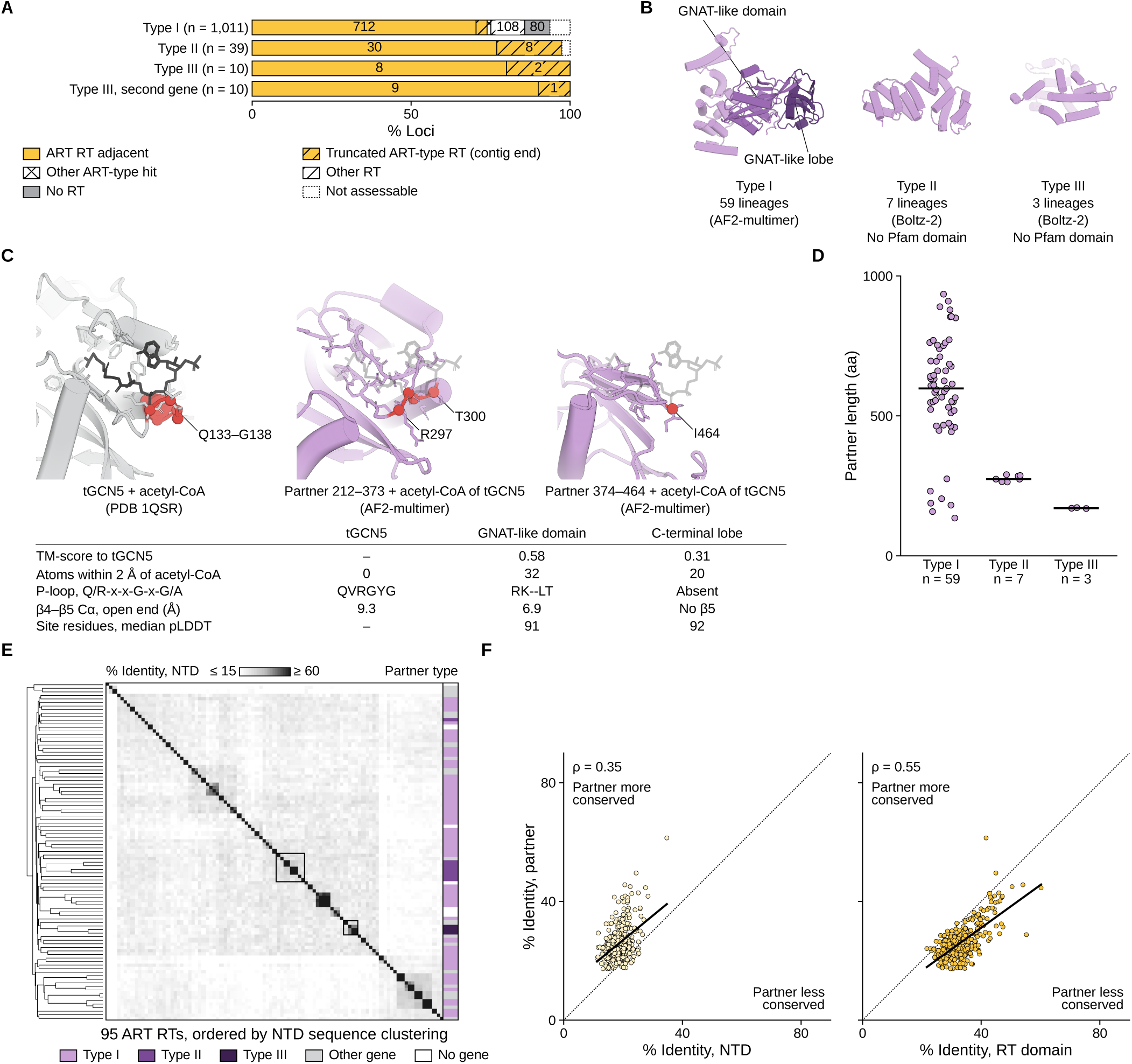
ART partner types and the RT N-terminal domain (NTD). (**A**) Database-wide occurrences of each partner family, by genomic context. (**B**) Predicted structure of one partner of each type. (**C**) Acetyl-CoA site of the GNAT acetyltransferase tGCN5 (PDB 1QSR; gray) compared to GNAT-like structures of ART partner proteins. (**D**) Partner length by type. Bars, medians. (**E**) Pairwise sequence identity of the NTDs of the 95 ART family RTs, ordered by hierarchical clustering. (**F**) Identity of the partner against identity of the NTD (left) and of the RT domain (right) for all 379 pairs of RTs that share a partner type.

## Acknowledgments

We thank F. Zhang, B. Hie, D. Li, and G. Brixi for reviewing an early copy of the manuscript and providing feedback. We thank the Anthropic life sciences team and the broader Anthropic community for building the models and tools that enabled this study.

## Materials and Methods

### Autonomous research harness

The campaign was run with an autonomous research harness in which every agent was an instance of Claude Code that was configured with Claude Mythos 5. The harness permitted up to 58 concurrent sessions, and agents ran in a sandbox with 60 CPU cores, 192 GiB of memory, and no GPU. A launch agent encoded the five stages of the research brief (Supplementary Note 1) as a chain, with each stage closed by scripted completion checks. The stages were 1) input assembly, 2) database sweep, 3) RT classification, 4) neighborhood census, and 5) deep dives. No task of a later stage was opened until the preceding stage was completed. Each task was defined by a task brief and assigned to a worker agent, which wrote and executed a plan before submitting a written summary of the results with the code and data files. A supervisor agent reviewed the plan, the summary, and the edited files, and either accepted them or returned them for revision. Of the 119 tasks, 49 were revised at least once. Follow-up tasks proposed by agents during the work entered a triage queue, from which the harness released or rejected with a written reason. After each completed task, a curator agent read the task brief and the worker’s summary and entered the findings into a shared knowledge base. Relevant entries were included in the prompts of later workers and supervisors. Reports were reviewed by an editor agent before filing. Plans, results, reviews, and scripts were committed to a version-controlled record readable by every agent. A campaign ended when no task remained that could be dispatched.

### Tools and skills available to the agents

Agents acted through the tools of Claude Code (a shell, file access, and web search). The harness added connectors to a metagenomic protein database, to literature search and full-text retrieval services, to the shared knowledge base, and to a queue for GPU jobs. Structure predictions were dispatched to NVIDIA A100 and L4 GPUs (19 jobs) and used either ESMFold (esmfold_v1) (62) or ColabFold 1.6.1 with AlphaFold2-ptm models (63). The sandbox provided Python with Biopython, NumPy, pandas, SciPy, and PyMOL, and agents installed further software as required. For sequence and RNA analysis, the agents ran HMMER 3.3.2 (64), MMseqs2 release 17-b804f (65), BLAST+ 2.16.0 (66), MAFFT 7.490 (67), FastTree 2.1.11 (68), SeqKit 2.8.2 (69), skani 0.3.2 (70), geNomad 1.12.0 (71), Infernal 1.1.5 (72), ViennaRNA 2.7.2 (73), and edlib 1.3.9 (74). For structure analysis, they ran TM-align (75) through tmtools 0.3.0 and the Foldseek web server (76). They also queried NCBI, UniProt, InterPro, the European Nucleotide Archive (ENA), the AlphaFold Protein Structure Database, the Protein Data Bank, and Rfam. In the retained session logs the agents issued 7,578 shell commands, 696 database queries, 131 literature searches, and 61 web requests. The sandbox also held a library of about 140 skills (written method guides) that workers consulted 15. Seven of these were guides for metagenomic surveys. They covered 1) counting the independent occurrences of a gene family, 2) scoring operonic arrangement against the background of the window, 3) adjudicating candidate families, 4) testing novelty claims, 5) annotating uncharacterized protein families, 6) proposing mechanisms together with the counterfactual evidence, and 7) drawing gene-neighborhood figures. The other libraries covered database use, literature access, GPU job dispatch, report editing, and the use of bioinformatic tools.

### Metagenomic sequence database

The database used in this study held predicted proteins from public metagenomic and genomic assemblies that spanned approximately 11 million biosamples, mainly from Logan (4), ENA, the Joint Genome Institute (JGI), and NCBI. Genes were predicted with prodigal-gv 2.10.0 (71, 77) under default parameters in unannotated assemblies of Logan, ENA, and NCBI, and the source gene calls were retained for annotated assemblies of NCBI and JGI. Each protein was linked to its contig, coordinates, strand, and biosample. Proteins were deduplicated, and were excluded if they had incomplete coding sequences, non-standard amino acids, more than 8,000 residues, degenerate k-mer repeats, or at least 50% low-complexity sequence as masked by tantan (78). The remaining proteins (approximately 15.8 billion) were clustered with DIAMOND (79, 80). Linear-time clustering was applied at 90% and then 70% identity with at least 80% coverage of the shorter sequence and was followed by cascaded clustering at 50% identity with at least 80% mutual coverage. Each cluster was represented by the member that was closest to the 80th percentile of length in the cluster. This yielded 1,939,242,578 clusters at 50% identity, whose representatives (hereafter, database representatives) formed the search space of the campaign and are the protein clusters that are counted in the main text. Agents accessed the database through a programmatic interface that returned the members of a cluster, protein and nucleotide sequences, gene neighborhoods, biosample records, and precomputed Pfam-A annotations (81). The interface also ran MMseqs2 searches (65) either against the database representatives or against a reference set of 365 million proteins (UniProt and NCBI nr, clustered at 90% identity).

### RT census by the agents

52 profile hidden Markov models (HMMs) were assembled from six Pfam profiles (RVT_1, RVT_2, RVT_3, RVT_N, GIIM, and Intron_maturas2) (81), the 45 class-specific profiles of myRT (82), and one profile that was built from an alignment from Toro (83). All database representatives were searched with hmmsearch (--noali -Z 8 --domZ 8 -E 0.01 --domE 0.01), which identified 3,391,714 candidate RT sequences. Candidates were retained if the RT core was complete (profile coverage ≥ 0.75) and the length reached a class-specific minimum (225 to 346 residues). The remainder were discarded as fragments (n = 2,016,377) or weak hits (n = 1,171,334; bit score < 25, coverage < 0.35, and no YxDD catalytic motif), or excluded for other reasons (n = 622). These selection thresholds were fixed beforehand on the five reference RTs that were named in the brief (Ec86, LtrA, BPP-1 Brt, AbiK, and an RT-Cas1 fusion) and on 2,339 labeled myRT sequences, which retained 99.0% of full-length labeled RTs. The 203,381 retained sequences were clustered with MMseqs2 easy-cluster (--min-seq-id 0.5 -c 0.8 --cov-mode 0) into 198,290 RT clusters. Clusters were then classified in three tiers. The first tier used 38 lineage-specific HMMs that were built from myRT (mafft --auto, hmmbuild) (67), with a class-specific bit-score floor and a margin of at least 10 bits over every other class (precision 0.997 in five-fold cross-validation). The second used MMseqs2 searches against labeled RTs (bit score ≥ 150, query coverage ≥ 0.5, identity ≥ 30%). The third used a FastTree tree (-lg -gamma) (68) of 2,241 RT domains that were aligned to RVT_1, which required agreement among the five nearest labeled leaves. This placed 35,168 clusters in the seven known classes, namely retron (11,517), unknown group (UG; 10,759), group II intron (5,461), diversity-generating retroelement (DGR; 3,857), CRISPR-associated (2,067), group II-like (827), and abortive-infection (Abi; 680). A further 25,737 clusters formed a loosely organized group on the tree apart from any labeled member and were treated as an eighth class of “novel” RTs. The 137,385 scattered clusters that could not be placed formed the ninth class of “unplaced” RTs.

### Sampling of RT neighborhoods

The worker that was assigned to the census defined a sampling cap for each RT class and drew 7,308 RT clusters as anchors. The abortive-infection (n = 680) and group II-like (n = 827) classes were sampled exhaustively, and the DGR and CRISPR-associated classes were sampled at random (1,000 clusters each). The UG class (700 clusters) was sampled in proportion to clade size, with clades of fewer than 30 clusters taken whole. Every retron cluster and every group II intron cluster carried a single clade label, so these two classes were sampled at random (1,000 retron clusters, with one control cluster added, and 500 group II intron clusters). The novel class was sampled at 60 clusters from each of its 16 clades, and the unplaced class was sampled with one representative of each of its 440 largest lineages plus 200 random singletons. At these caps, the worker estimated that a family had to occur at 0.3 to 0.6% of the loci of a class to reach three independent occurrences. For each anchor, up to three loci were retrieved from each of four source collections (Logan, ENA, JGI, and NCBI) with 10 kb of flanking sequence on each side. Loci were recovered for 7,238 of the 7,308 anchors (10,983 loci with 79,680 coding sequences), and 85.9% of the windows were truncated by a contig end.

### Scoring of partner families

Neighboring proteins were grouped into families either by their best Pfam-A match or, where none was found, by MMseqs2 clustering. The 3,564 families that recurred across loci were evaluated against three criteria (promotion filters). The worker that was assigned to the census had set the thresholds of these criteria before the census was run. First, the family had to be conserved within at least one RT clade. Second, its occurrences had to be independent, meaning that they were found in at least three clusters at 90% identity, in at least three biosamples, and beside RTs of at least two different classes. For example, families that were found only beside retrons could not meet this criterion, because all retrons carried one clade label. Third, genes of the family had to lie nearer to the RT than randomly drawn genes of the same loci (P ≤ 0.05). They also had to lie either on the strand of the RT or within 100 bp of the adjacent gene. Known partners of RT systems and general residents of mobile elements were designated as controls beforehand. Forty-six families met all three criteria, and 13 of these were controls that included Avd, Cas1, and Cas2. Of the other 33, the worker promoted 16 after a review of their annotations. A seventeenth family was promoted later by a follow-up task.

### Deep dives

In each deep dive, the worker re-applied the three criteria to the members of the promoted family, examined alignments, and searched annotations and the literature for prior descriptions of the association. A family was counted as confirmed when its report upheld it as a new RT partner. Otherwise it was counted as rejected or set aside.

### Campaign accounting

The research campaign comprised 119 tasks. Of these, 107 were completed, 10 were rejected at triage, and 2 were stalled (one after ten revisions and one after ten failed completion checks). One task assembled the input data, 96 performed analyses, and 22 wrote or revised a report. Five tasks were seeded from the stages of the brief, 16 were deep dives that were seeded from the promoted families, and 98 were follow-up tasks. The agents ran 949 sessions (the launch session and 414 worker, 375 supervisor, 107 curator, and 52 editor sessions). Agent-hours were calculated as the summed duration of all agent sessions, including the execution of tools within sessions (76.9 agent-hours, 63.7 of them in worker sessions). Token use was calculated as the sum over all sessions of uncached input tokens (11.3 million), output tokens (14.9 million), and tokens that were written to the prompt cache (189.5 million), which gave 215.6 million tokens. Tokens that were read from the prompt cache were excluded. To attribute token use to the stages of the campaign, the tokens of each session were assigned to its task, and each follow-up task was assigned to the survey stage or deep dive from which it descended in the task record. To determine whether each follow-up task originated with a worker or a supervisor, a Claude instance was given the brief and worker summary of the parent task and the brief of the follow-up task. It classified each follow-up task as either proposed by the worker or originating in the supervisor’s review. The classification was run twice, and the six discordant calls were resolved by reading the task records.

### Ranking of reports

The 19 reports of the campaign were ranked in a tournament of pairwise comparisons, a design that was used previously to rank machine-generated hypotheses (21, 24, 84). Every ordered pair of reports was compared once (342 games). In each game, a language-model judge (Mythos 5) read a fixed rubric and two reports under anonymized file names before scoring each report from 1 to 5 for impact, novelty, soundness, and actionability (weights 0.35, 0.30, 0.25, and 0.10). The report with the higher weighted score won, and a report with a soundness score of 2 or less lost automatically. Ratings were fitted to all games with a Bradley–Terry model. The ART report ranked third (32 wins in 36 games).

### Transcript analysis of the discovery

Claude Code records each session as a log of reasoning blocks, messages, tool calls, and tool results, which along with messages make up the reasoning trace. The log of task t0062, in which a worker recognized the ART array, was parsed into an ordered list of events. Recognition was placed at the first reasoning block or message that described a repeat array, and the tool results just before it were taken as what the worker had read. The brief of this task, which was written by the supervisor agent of the preceding task, requested a test for a retron-type ncRNA upstream of the RT. Neither this task brief nor the research brief mentioned repeats or arrays. The two retrievals before recognition returned sequence without annotation, and no tool call came between them and recognition. No earlier tool call contained a repeat analysis, and no established repeat finder was run in the session. Every repeat that the worker named was recounted in the retrieved sequences. The first repeat pattern that the worker wrote (CATGTGT[AT]TCGCATGT) was recounted on both strands of the 2,900-nt sequence that the worker had retrieved, with at most 1 mismatch. This sequence is the flank of locus L0050, the worker’s label for the locus at which it first recognized an array. Quotations from the session logs are verbatim, including the agents’ emphasis marks, with omissions marked by bracketed ellipses and shading added. Loci that the agents examined were drawn to scale from the gene tables that were printed in the logs. Repeat positions were obtained by re-scanning the logged sequences with the worker’s repeat core, with at most 2 mismatches allowed. For phage MarsHill, the GenBank record (MW248466.1) was scanned.

### Analyses that the agents ran on the array

The agents assembled the RT family from MMseqs2 searches and took as members the RTs that were linked to one another by hits above a score threshold. In each upstream region the most frequent 14-mer was taken as the repeat, its copies were located with up to two mismatches, and the sequences between neighboring copies were taken as spacers. For each of the 42 spacers from six arrays, the longest stop-free peptide was searched against a reference set of 365 million proteins. The spacers of two cultured phages were each aligned to the genome of the other phage with edlib. The folding energy of 40 spacers was predicted with ViennaRNA and compared with that of 30 shuffled sequences of the same dinucleotide composition. Novelty was tested against the annotations of family members and the literature on retrons, defense-associated RTs, AbiK, and the CRISPR arrays of jumbo phages. The fractions that are given for these analyses (spacers without a protein match at E ≤ 0.001, and spacers with z < −2 against the shuffled sequences) were taken from the agents’ report and were not recomputed.

### Human-directed analyses

All analyses that were performed after the campaign were carried out in interactive Claude Science sessions, in which the authors directed the analysis and Claude wrote and ran the code.

### Family definition

Homologs of the MarsHill RT (GenBank QQM14740.1) were collected from GenBank with BLASTP (66) and from our metagenomic database with a profile HMM. The profile was built from 12 of the GenBank phage RTs (MAFFT --auto, hmmbuild). Ten of these came from *Staphylococcus* phages and two from *Listeria* phage LPJP1. The profile was searched against all database representatives with hmmsearch (HMMER 3.4; E ≤ 1 × 10^−5^), which returned 9,100 proteins. Hits of 350 to 900 residues (n = 4,379) were pooled with the 12 phage RTs and 2,019 reference RTs of the myRT classification (82). The pooled set was compared all against all with DIAMOND blastp (--very-sensitive, E ≤ 1 × 10^−5^) (79), and proteins were linked at bit scores of at least 140. The connected component that contained the MarsHill RT held 117 proteins and no reference RT. All members of the database clusters that were represented in this component were then retrieved, and the 706 further members of at least 400 residues were added to the 117. The resulting 823 full-length RTs were clustered with MMseqs2 easy-cluster (--min-seq-id 0.9 -c 0.8) into 230 clusters, and the member on the longest contig was taken as the representative of each cluster. Representatives were aligned with FAMSA (85), and a tree was inferred with FastTree under the LG model (68). The ART family was defined as the smallest clade of this tree that contains every locus with both a type I partner gene and a non-coding region of at least 500 nt upstream of the RT. Partner types are defined under Partner genes and partner types. This clade held 95 representatives, which were numbered ART_01 to ART_95 in tree order. Members that were encoded on GenBank phage genomes were classed as cultured phages. Contigs were classified as virus, plasmid, or chromosome with geNomad 1.12.0 (database release 1.9, end-to-end workflow, default settings) (71), and gene calls on database contigs were taken from the database annotation.

### Repeat-array detection

Arrays upstream of RTs were handled in three steps, with each step described under its own heading. They were 1) detected with a k-mer scan, 2) delimited and measured with a procedure that does not fix the repeat length, and 3) annotated copy by copy for sequence comparisons between arrays. The scan searched up to 3,000 nt upstream of the RT start codon for short words that recurred at least three times at regular spacing (100 to 450 nt, start to start) and was run with two word settings. For the 95 members, the 20 most frequent 14-nt words of the window served as seeds (at least three distinct bases, no homopolymer of six or more). Copies were non-overlapping windows within two mismatches of a seed. An array was called when the longest run of regularly spaced copies, R, was at least 3 and exceeded the longest such run in 100 mononucleotide shuffles of the window. No array was called when a longer run of copies that were spaced under 100 nt apart was present. R was taken as the copy number of the array. One locus with R = 3 that did not exceed its shuffles was retained because an exact 12-nt word recurred three times at such spacing. For the tips of the RT phylogeny, which include retron loci, an array was called either from annotated repeat copies (see Array annotation and comparison) or from an exact 12-nt word that recurred three times at such spacing. Under both settings, loci with less than 1,500 nt of contig upstream of the RT and no array were recorded as not assessed. Neither setting called an array at any NART locus.

### Array delimitation and measurement

Up to 6,000 nt upstream of the RT was rescanned in the 95 members and in 8 further GenBank phage ART loci with every recurring 10-nt word as a seed. Copies were collected at up to one mismatch per seed. The longest chain of copies at near-constant spacing (60 to 600 nt, 30% tolerance, single skipped copies allowed) was retained for scoring. The score of a chain was the number of copies minus one, multiplied by the information content of its conserved block relative to the base composition of the region. A chain was retained when its score exceeded the best chain score in each of 200 shuffles of the region, in both a 3,000-nt and a 6,000-nt window. Shuffles permuted nucleotides within consecutive 50-nt blocks to preserve local base composition. Chains that scored less than twice the best shuffle score were retested against 2,000 shuffles. A position weight matrix of the repeat was then built, and matches that scored above the maximum of 200 shuffled regions were counted as copies. The repeat was taken as the contiguous run of columns in which at least 80% of copies carry the consensus base, with one lapse tolerated. The repeat length was measured both on ungapped copies and on a MAFFT alignment of the copies. Chains whose spacings were all multiples of three and whose copies lay within an annotated gene were set aside as coding repeats. An array was called adjacent to the RT when no annotated gene of 300 nt or more lay between its last copy and the RT. Each locus was also scanned with the position weight matrix of every other array to group arrays that share a repeat. Copy numbers, spacings, repeat and spacer lengths, and array spans that are given in the text are from this procedure, which delimited arrays adjacent to the RT at 28 of the 95 loci and detected none at NART loci.

### Reference RT sets

Two reference sets of RTs were assembled. The 957-sequence set, which was used to compare N-terminal lengths across RT classes, comprised 633 labeled RTs from published classifications (39, 43, 82), 294 classified representatives from our database, and 30 ART RTs. Each lineage of the published classifications was clustered at 50% identity with MMseqs2. Labeled references were then sampled with a quota per lineage that was proportional to the square root of its size. The 294 database representatives were drawn from one of 64 equal partitions of the database. They had been assigned to a class by their best-scoring profile in a panel of RT class profiles (E ≤ 1 × 10^−10^, profile coverage ≥ 0.60) and were clustered at 40% identity. The 774-sequence set, which was used for the RT phylogeny, comprised 98 ART and NART RTs, 617 retron RTs, and 59 group II intron, DGR, UG/Abi, and CRISPR-associated RTs. Of the retron RTs, 325 came from published classifications (39, 82) and 292 were retron-classified representatives from our database. The published retron RTs spanned retron clades 1 to 11 and the retron-like RTs of these classifications, and included the 13 references that were most similar to MarsHill RT homologs in a DIAMOND search (79).

### Phylogeny

RT-domain sequences of the 774-sequence set (taken as the Pfam RVT_1 envelope (81) extended by 40 residues on each side) were aligned with MAFFT L-INS-i (67) and trimmed with trimAl (-gappyout) (86), which left 254 columns. A maximum-likelihood tree was inferred with IQ-TREE 3.1.2 (87) under the model that ModelFinder (88) selected by BIC (Q.pfam+F+R6), with 1,000 ultrafast bootstrap (89) and 1,000 SH-aLRT (90) replicates, and was midpoint-rooted. In this tree the ART and NART RTs form the two child clades of the node that unites them (71 and 29 tips, which include two database representatives that had been sampled as retrons). For display, every maximal clade with at least 90% of tips from one RT class was collapsed.

### N-terminal region

The N-terminal region of an RT was defined as the residues that precede the envelope of its first Pfam RVT_1 (PF00078) domain (hmmsearch, HMMER 3.4; independent E ≤ 1 × 10^−3^). This boundary was used throughout except in Fig. 2D and Supp. Figs. 3A, 5A, and 5E, where the region ends 40 residues before the envelope, at the start of the RT-domain sequence that was used for phylogeny. N-terminal lengths were compared across the classes of the 957-sequence set (group II intron and group II-like, n = 226; CRISPR-associated, 125; DGR, 32; UG or Abi, 434; retron, 110; ART, 30).

### Catalytic-motif logos

The YxDD motif was located with the pattern [YF].DD in the 95 ART members and in RVT_1 domain sequences from the 957-sequence set. These domain sequences came from the retron (n = 110), DGR (n = 32), and group II intron (n = 206) representatives. Windows of 10 residues on either side of the motif were stacked without gaps. Letter heights are residue frequency multiplied by column information content (log_2_20 minus Shannon entropy).

### Structure prediction

AlphaFold2 (91) models were built with ColabFold 1.6.1 (63) for the 95 ART members, the RTs of the 8 further GenBank phage ART loci, 29 NART RTs, and 32 retron RTs that are closely related to them. Settings were alphafold2_ptm, 5 models, 3 recycles, no templates, and no relaxation, and the top-ranked model was used. Each MSA combined the ColabFold MMseqs2 server alignment (UniRef30 2302, ColabFold environmental database 202108) with an alignment of related RTs that was mapped onto the query sequence. The MarsHill RT model that was used for the structural comparisons was built with the same ColabFold version from the server alignment alone (3 models, 3 recycles; top-ranked model used).

### Structural comparisons

Retron Ec86 (PDB 7V9X, chain B), DGR BPP-1 (PDB 8UBE, chain A), and the NART model were superposed on the MarsHill model over the RT domain in PyMOL. N-terminal residues with a per-residue model confidence (pLDDT) below 50 were not drawn. The RVT_1 ranges of Ec86, BPP-1, and the GsI-IIC group II intron RT (PDB 6AR1, chain A) were superposed on the MarsHill RT domain with TM-align (75). Structural similarity was reported as the TM-score that was normalized by the length of the MarsHill RT domain. The MarsHill RT core (residues 211 to 501) and N-terminal region (residues 1 to 210) were each searched against a panel of 16 solved RT structures with Foldseek 10.941cd33 (easy-search--exhaustive-search 1 -e 10) (76). The panel comprised PDB 7V9U, 8QBK, 8QBL, 8QBM, 9NNB, 9P4J, 8UB9, 5HHL, 6AR1, 9C0I, 9IOA, 7R06, 7R07, 7R08, 8OZ7, and 1RTD. It was chosen by hand to cover solved RTs of retrons, a DGR, group II introns, DRT2, DRT9, and abortive-infection systems, with HIV-1 RT as an outgroup. The highest query-normalized TM-score over the chains of an entry was taken as the score of that entry. To cluster N-terminal folds, N-terminal regions of at least 60 residues with mean pLDDT of at least 50 were compared all against all with TM-align on N-terminal coordinates only (132 of 164 models; 87 ART, 29 NART, 16 retron). The TM-score that was normalized by the longer region was used. Structural clusters were defined by average linkage on 1 − TM-score, cut at 0.5.

### N-terminal sequence comparisons

To compare the divergence of N-terminal regions with that of RT domains, each locus was paired with the locus whose RT domain was most similar. Identities of the RT domains and of the N-terminal regions were computed from global alignments (BLOSUM62, gap open −11, extension −1) as identical positions over the length of the longer sequence.

### Locus features by clade

Four locus features were recorded for each tip of the ART (n = 71), NART (n = 29), and nearest retron (n = 13) clades of the RT phylogeny and were counted only where they could be assessed. Repeat arrays were called as described under Repeat-array detection. A non-coding tract was recorded when at least 500 nt separated the RT from the preceding gene. An ART partner was recorded when the protein that was encoded directly downstream on the RT strand matched the partner protein of an ART locus (phmmer, E ≤ 1 × 10^−3^). An RNA polymerase was recorded when an upstream gene on the RT strand carried a Pfam RNA-polymerase domain.

### Comparison of the SA1 and MarsHill loci

Loci were taken from the GenBank records of phages MarsHill (MW248466.1) (61) and SA1 (MW218148.1) (48). Repeat copies were located as matches to the 14-nt core ATATGAATACGTAT with at most one mismatch. The MarsHill array region, from 60 nt before the first core to the RT start codon, was aligned globally to that of SA1 (match 2, mismatch −1, gap open −4, extension −0.5). The RT genes were aligned as codons, with a global protein alignment (BLOSUM62) as the guide. Per-nucleotide identity was smoothed with a centered 25-nt running mean. The five MarsHill copies were aligned without gaps at the core with 29 nt on either side, and positions with the same base in at least four copies were classed as conserved.

### Array annotation and comparison

For sequence comparisons between arrays, repeat copies were annotated in ART loci that hold at least three exact copies of a locus-specific 10-nt repeat word, with each copy within 600 nt of the next. The word of each locus was taken from a scan of up to 3,000 nt upstream of the RT for 10-nt words that recurred in tandem (at least three exact copies). The periodicity of the word was 141 to 317 nt in the 24 loci with such a word. A 26-nt consensus (6 nt before to 20 nt after the start of the word) was built from the exact copies. The strand was then rescanned, and windows that matched at 21 or more of 26 positions were taken as copies. Gaps of at least 1.6 times the usual spacing of an array were then searched, within 40 nt of each expected position, for degenerate copies that matched at 18 to 20 positions. Of the 24 annotated arrays, 16 were analyzed. Five *Staphylococcus* phage arrays that are closely related to those of MarsHill and SA1 were omitted, and three loci were excluded (no usual spacing definable, too few exact copies, or RT outside the ART clade). To profile conservation along the array unit, windows from 60 nt before to 160 nt after the start of each exact copy were stacked without gaps. Identity at each position was the mean over all pairs of copies and is 100% at the 10-nt word by construction. To compare repeats, flanks, and spacers between arrays, each copy was divided into the 26-nt repeat, its flanks (20 nt on either side), and the spacer between the flanks of consecutive copies. Repeats and flanks were compared without gaps, and spacers were aligned globally (match 1, mismatch −1, gap open −3, extension −1). Chance identity was the same measure with one sequence of each pair shuffled (median of 20 mononucleotide shuffles).

### RNA-seq analysis of published SA1 infection data

RNA-seq reads of SA1 infection of *Staphylococcus lentus* (BioProject PRJNA836150 (48); 12 paired-end, strand-specific libraries from three cultures sampled before infection and at 5, 15, and 55 min after infection) were downloaded from ENA. Reads were trimmed with fastp 1.3.6 (minimum length 30 nt) (92) and aligned with Bowtie 2 v2.5.5 (--very-sensitive -X 1000 -- no-unal) (93) to the SA1 genome (MW218148.1) together with the *S. lentus* chromosome (NZ_CP059679.1). Alignments were sorted and indexed with SAMtools 1.24 (94). Properly paired alignments with MAPQ of at least 10 and a template of at most 1,500 nt were retained as fragments. Each fragment was counted once on the strand of read 2 (the sense read) and assigned to the feature that contained its midpoint. Features were the 258 annotated SA1 coding sequences and the array RNA, which was defined as the whole non-coding region upstream of the RT (positions 9,448 to 10,646). Transcripts per million (TPM) were computed from sense fragments per kilobase over these 259 features. TPM was averaged over the three cultures of a time point, and features were ranked by this mean. The share of phage RNA was calculated as the number of array-RNA fragments divided by all fragments that aligned to the SA1 genome in that library. Coverage was calculated as the number of fragments that spanned each position per million phage fragments. Array coverage was averaged over the three 15-min cultures and, for comparison of replicates, was scaled to the maximum of each library over the array.

### ART expression plasmids and cultures

Two plasmids that carry parts of the SA1 ART locus (GenBank MW218148.1) were assembled by Golden Gate cloning and verified by whole-plasmid sequencing. The native-locus plasmid carries positions 9,448 to 12,951 (non-coding region, RT, and partner genes). This insert was synthesized as five fragments and assembled downstream of a P_LtetO-1_ promoter on a pSC101-origin vector that encodes TetR and ampicillin resistance. The clone that was used carries one substitution in the non-coding region (C to T at position 9,936). The heterologous-promoter plasmid carries the 1,187-nt non-coding region behind the J23119 promoter and a synthesized RT gene behind the *araBAD* promoter, on a pMB1-origin vector that encodes AraC and kanamycin resistance. Its RT gene was recoded to encode the 505-residue protein that starts at position 10,635, 12 nt upstream of the annotated start codon, and its non-coding region was amplified by PCR from a native-locus clone. Plasmids were carried in *E. coli* DH10B, and one culture per construct was grown at 37 °C in Terrific Broth with 100 µg/mL carbenicillin (native-locus plasmid) or 50 µg/mL kanamycin (heterologous-promoter plasmid). The native-locus culture was grown without anhydrotetracycline to an OD_600_ of about 0.4 and harvested 2 h later. The heterologous-promoter culture was diluted 20-fold from an overnight culture into medium with 0.2% L-arabinose and harvested after about 4 h.

### Small RNA sequencing

RNA was extracted with hot formamide as previously described (95, 96). In brief, cell pellets were resuspended in 95% formamide with 18 mM EDTA at 65 °C for 5 min. The cleared extract was purified on RNA Clean & Concentrator-5 columns (Zymo Research), which retain RNAs of 17 nt and longer, with in-column DNase I digestion. No size selection was applied. Ribosomal RNA was depleted from 500 ng of RNA with the NEBNext rRNA Depletion Kit (Bacteria) (New England Biolabs). RNA ends were repaired with Quick CIP and then with T4 polynucleotide kinase, first without and then with ATP (New England Biolabs), and RNA was recovered on SPRI beads. Libraries were prepared with the NEBNext Multiplex Small RNA Library Prep Set for Illumina (New England Biolabs) following the manufacturer’s protocol, except that adaptors and the reverse-transcription primer were diluted twofold. Libraries were cleaned with 1.5 volumes of SPRI beads. Pooled libraries were sequenced on a NextSeq 2000 (Illumina) with 2 × 151-nt reads and 20% PhiX. Reads were demultiplexed and adapter-trimmed with BCL Convert 4.2.11 (Illumina) and filtered with fastp 1.3.6 (adapter trimming off, poly-G trimming, minimum length 12 nt). Filtered reads were aligned with Bowtie 2 v2.5.5 (--end-to-end -D 20 -R 3 -N 0 -L 12 -i S,1,0.50 -X 1000) to the plasmid of each library together with the *E. coli* DH10B genome (NC_010473.1) and phiX174. Alignments were sorted with SAMtools 1.24 (94). Fragments were taken from the start of read 1 to the end of read 2 for proper pairs and from the read itself otherwise, with read 1 in the sense orientation. Secondary and supplementary alignments, fragments longer than 1,000 nt, and alignments with MAPQ below 2 were discarded. Coverage was reported as raw sense-strand fragment counts in SA1 genome coordinates.

### Array RNA boundaries

Array RNAs were called as contiguous runs of at least 40 nt at or above half of the maximum coverage over the array. RNA 1 was called in the *E. coli* library that expressed the array from the J23119 promoter, and RNAs 2 and 3 were called on the mean coverage of the three 15-min infection cultures.

### RNA secondary-structure prediction

Minimum-free-energy structures were predicted with RNAfold of ViennaRNA 2.7.2 (73) at 37 °C with default parameters and drawn at one scale within a panel. The Ec86 ncRNA was the 169-nt interval between the outer inverted repeats of retron Ec86 (M24408.1, positions 373 to 541), and SA1 RNAs 1 to 3 were the intervals that were called from coverage. Array units were cut from the arrays of seven *Staphylococcus* phages, SA1 (MW218148.1), MarsHill (MW248466.1), Madawaska (MW349129.1), LY01 (OR836606.1), S6 (LC680885.1), PALS_2 (MN091626.1), and UFV_DC4 (MZ779063.1). A repeat was defined as the 22 nt from 4 nt before to 17 nt after the start of the 14-nt core, and a spacer was defined as the sequence between consecutive repeats. Units of the SA1 array were built from repeats 3 and 4 and spacers 3 and 4. For each of the seven arrays, two units were folded. One was formed by repeat 3, spacer 3, and repeat 4, and the other was formed by repeat 3 and spacer 3.

### Partner genes and partner types

The partner of each of the 95 ART RTs was taken as the first gene downstream of the RT. Proteins that were encoded in the 230 representative loci were clustered with MMseqs2 cluster (--cluster-mode 1 -e 0.01 -s 7.5 -c 0.5), and the cluster alignments were compared all against all with HHblits (2 iterations, E ≤ 0.01) (97). Clusters whose profiles matched with at least 90% probability over at least half of both profiles were merged into families. The three families that recurred as partners of ART RTs were designated types I to III, and type III loci carry a second recurring family directly downstream of the partner. Similarity between types was assessed on predicted structures of eight partners (six of type II, one of type I, and one of type III; see RT-partner co-folding and partner models). The eight were compared all against all both by sequence (phmmer, HMMER 3.4) and by structure (TM-align, after removal of residues with pLDDT ≤ 50). Members of each family were also searched against the profile HMMs of the other families. Thirty-six partner sequences of the three types were searched against Pfam-A (81) with hmmsearch at gathering thresholds (--cut_ga), with the search-space size set to the number of models.

### Database-wide occurrence of partner families

A profile HMM was built for each partner family (MAFFT L-INS-i, hmmbuild) from 46 type I partners, 13 type II partners, and the 8 and 6 database members of the two type III genes. Each profile was searched with hmmsearch against all database representatives. A representative was assigned to a family when it scored at or above the lowest-scoring known member (leave-one-out score for families of at least 10 members, self-score otherwise). The cluster of each assigned representative was expanded to all member proteins, and every locus was retrieved as the partner gene ±6 kb (1,011 type I and 39 type II loci, and 10 loci for each type III gene). Annotated proteins and six-frame ORFs of at least 40 residues in each window were searched against a panel of 221 RT profiles (hmmsearch, E ≤ 1 × 10^−5^). Loci were then assigned to the first matching category of six. An ART RT was called adjacent when an RT that scored at least 207 bits on the ART profile was co-oriented with the partner and lay one to three genes upstream. A truncated ART-type RT was such an RT that scored below the threshold and abutted the contig end. The remaining categories were another ART-type hit, another RT, no RT (no hit and at least 1,800 nt of contig upstream of the partner), and not assessable (no hit and less than 1,800 nt upstream).

### RT-partner co-folding and partner models

Type I RT-partner pairs from five environmental loci and one shuffled non-cognate pair were co-folded with AlphaFold2-multimer v3 (98) in ColabFold 1.6.1 on locus-paired MSAs (3 models per pair, 3 recycles, 1 seed, no templates). Each MSA opened with a paired block. To build this block, the RT and partner sequences of type I loci whose partner starts within 60 nt of the RT stop were aligned to the query pair (MAFFT --auto) and joined by locus. Unpaired blocks of MarsHill RT homologs and of type I partners followed (MAFFT --add --keeplength), and no server alignment was used. In the shuffled pair, the paired rows joined RTs and partners of different loci. Type II pairs of phages SA1, MarsHill, LY01, and PB50 and of two metagenomic loci were co-folded with two engines. AlphaFold2-multimer v3 was run in ColabFold 1.6.1 (5 models × 3 seeds, 3 recycles, no templates), and Boltz-2 v2.2.1 (99) was run with 3 recycling steps and 5 diffusion samples × 2 seeds. One type I pair served as a positive control, and the SA1 RT with a type I partner served as a negative control. Both engines received the same MSAs. These MSAs combined 1) a paired block of same-locus RT and partner sequences from the other loci of that partner type, 2) unpaired ART family RT and partner sequences, and 3) ColabFold MMseqs2 server hits (UniRef30 2302, environmental database 202108). The interface confidence score (ipTM) is the value that each engine reported, which was summarized per pair and engine as the median over all models. The type I partner and complex displayed in the figures are from the top-ranked AlphaFold2-multimer model of the ART_57 pair, and the type II partner and complex are from a Boltz-2 model of the SA1 pair. The type III partner is the most confident Boltz-2 model of a co-fold of the two type III genes of one locus. Models were drawn with residues of pLDDT below 50 hidden, and for comparison with solved RT complexes the RT of each co-fold was superposed on the RT of PDB 8UBE.

### Comparison of the type I partner with a GNAT acetyltransferase

The ART_57 type I partner model was compared with *Tetrahymena* GCN5 (tGCN5), a GCN5-related N-acetyltransferase (GNAT), bound to acetyl-CoA (PDB 1QSR, chain A). GNAT-like regions were located by TM-align of tGCN5 against the whole chain and then against the chain with the first hit removed. The first hit defined the GNAT-like domain (residues 212 to 373), and the remaining C-terminal residues (374 to 464) were analyzed as a second region. Each region was superposed on tGCN5 with TM-align (TM-scores normalized by tGCN5), and the acetyl-CoA of 1QSR was kept at its crystallographic position. For each region, partner heavy atoms within 2.0 Å of acetyl-CoA were counted, and residues with any atom within 4.0 Å were recorded with their median pLDDT. The residues that aligned to the P-loop of tGCN5 (residues 133 to 138) were identified, and the Cα-Cα distance between strands β4 and β5 at their open end was measured. Strands were assigned on 1QSR with DSSP (100).

### N-terminal region and partner divergence

The N-terminal regions of the 95 ART RTs were aligned pairwise (global, BLOSUM62, gap open −10, extension −0.5). Identity was calculated as the number of identical aligned residues over the length of the longer sequence, and RTs were ordered by average-linkage clustering of 1 − identity. Partner divergence was related to RT divergence on a subset of ART tips of the RT phylogeny (30 RTs). These were the tips whose downstream gene lay on the same strand and was a type I, II, or III partner, and whose N-terminal region was at least 100 residues. For all 379 pairs of these RTs that share a partner type, the N-terminal regions, RT domains, and partners were aligned, and their identities were calculated in the same way. Spearman correlations and least-squares lines are shown, and no P value is given because pairs that share an RT are not independent.

### Replicate campaigns

The campaign was launched ten more times with the same harness and brief. All task records (n = 3,084), reports, and session transcripts (n = 5,632) of the replicates were searched for identifiers from the original campaign. These were the identifiers of the RTs (n = 130) and contigs (n = 171) that the agents of the original campaign had assigned to the RT family, and the names and accessions of cultured phages that encode such RTs. Sessions that named one of these contigs (n = 11) were parsed event by event for tool results that contained a contiguous DNA string of at least 200 nt and for remarks on repeats. An ART locus on a contig outside this identifier set would not be found by this search.

### Fixed-input benchmark

Each attempt was an independent run of one model that received the same task statement and one of five levels of input. The task statement asked the model to characterize the RT and to “predict its function: report every feature you judge important, the evidence for it, how confident you are, and what experiment would test your main prediction.” It stated that the loci had been drawn from a large sequence collection because each encodes a member of one related group of RTs, and that nothing else was known. Each attempt ended with a report and a structured submission of findings, in which each finding consisted of a claim, its evidence, and a confidence between 0 and 1. The inputs were 96 ART loci (1,093 to 26,554 bp) with anonymized identifiers, gene calls, and Pfam matches, and 1,131 predicted protein structures. At level 1 the prompt contained two RT protein sequences, and at level 2 it contained the two corresponding loci as text. Neither level provided tools. At level 3 no sequence was in the prompt, and all 96 loci were supplied as files in an offline environment with gene calling, HMMER, BLAST+, MMseqs2, MAFFT, FastTree, ViennaRNA, and Python. Level 4 added a predicted structure for every open reading frame, Foldseek with a local reference structure database, PyMOL, and US-align (101). Level 5 added literature search, web access, and the installation of software. The seven Claude models that are named in the Results were each run for 100 attempts at every level (3,500 attempts). Each attempt was run in isolation in Claude Code with a maximum output budget of 1 million tokens.

### Benchmark grading

Submissions were graded on the ten binary claims that are listed in Fig. 4B, which the authors selected as characteristic of the ART family. Three claims concerned the 5′ non-coding tract, two concerned the RT, and five concerned the partner. Grading was performed by a language-model judge (Mythos 5) that was presented only with the submission and the rubric. A programmatic keyword screen identified the findings that were relevant to each claim for the judge and did not contribute to the grade. The judge graded each claim independently. A claim was counted only when the submission asserted it as a conclusion and not as one of several hedged possibilities, and no credit was given for over-claims that were specified in the rubric. The score of an attempt was the number of asserted claims (0 to 10). An attempt was counted as having recognized the repeat array when the judge graded the repeat-array claim as asserted.

### Genomic language models

The 2,900-nt flank of locus L0050, as returned to the worker in task t0062, was evaluated with gLM2 (50) and Evo 2 (49). The gLM2 model (gLM2_650M) received the whole flank as one input. Each nucleotide was masked in turn, and the probability of the true base was taken from a softmax over the four nucleotides at the masked position. Evo 2 (7B model) was accessed through the Evo Designer web interface, and the probability of each base given the preceding sequence was taken from the returned logits, which were renormalized over A, C, G, and T. Values are the natural-log probability of the true base minus ln 0.25, so that zero corresponds to a uniform guess among the four bases. As a control, the nucleotides of each of the 14 copies (16 nt each), and of one degenerate copy upstream of the first, were permuted in place with the rest of the flank unchanged, and both models were run again. Tracks are centered 16-nt running means. Aligned profiles are means over copies 2 to 14 in 3-nt bins from 24 nt before to 41 nt after the start of the repeat, and per-copy values are means over the 16 nt of a copy.

### Mythos 5 internal signals during the discovery session

The transcript of the session of task t0062 was reconstructed from the session log, up to and including the reasoning block in which the model first reports the array, and passed once through a separately hosted copy of the same Claude Mythos 5 checkpoint (saved version of the model) that ran the session (Fig. 5, G to I). The worker’s system prompt and tool definitions are not in the log and were left out (placeholder tool definitions were used). The model’s activity at one layer (one of its successive processing stages) was decomposed with a sparse dictionary (102, 103): a set of recurring activity patterns learned without supervision from the model’s own activity, each with an automatic description generated from the texts on which it is strongest. An internal signal is the strength of one pattern at one token (the model’s unit of text). Only the 128 strongest signals are recorded at each token, so a zero means no strong response. The model reads text from left to right, and its activity at a token depends only on that token and the text before it, so a response at copy 3 cannot depend on later copies. Each nucleotide takes the value of the token that contains it, a copy’s value is the mean over its 16 nucleotides, and a nucleotide counts as responding if its token has a recorded value.

Candidate signals were selected before flank L0050 (Fig. 1F) was examined: the 12 with the largest differences among those that responded to synthetic repeats, that is, repeats planted in computer-generated random sequences (to qualify, a signal had to be higher on the third and later copies than on a twin sequence in which every copy was shuffled, and higher than between the copies, in at least 9 of 10 sequence pairs). The two shown were chosen from these 12 after the flank had been examined: the signal labeled automatically as “tokens repeating an earlier pattern/text verbatim” (repeat-signal 1), and the signal labeled as “letters in repeated DNA/protein sequence motifs, and garbled text” (repeat-signal 2). Two controls support the text. With each of the 14 copies shuffled in place in the same transcript, repeat-signal 2 was 0.00 on all 14 copies, against per-copy means of 0.85 to 1.62 on copies 3 to 14 in the flank as the model retrieved it. On repeats planted in random strings of letters and digits (10 pairs), repeat-signal 1 was 0.48 on the fourth and later copies against 0.05 on the shuffled twins, and repeat-signal 2 was 0.75 against 0.01 (each higher in 10 of 10 pairs); by the same measure repeat-signal 2 was 0.69 against 0.00 on 10 further computer-generated DNA pairs that played no part in the selection (the two sets differ in more than the alphabet, so the two values are not compared with each other). Both signals therefore respond to repeats that are not DNA; neither is specific to DNA. Tracks, aligned profiles, and per-copy values are displayed as in Fig. 5, D to F.

